# Protein stability engineering insights revealed by domain-wide comprehensive mutagenesis

**DOI:** 10.1101/484949

**Authors:** Alex Nisthal, Connie Y. Wang, Marie L. Ary, Stephen L. Mayo

**Author notes:** Present address: Xencor Inc., 111 W. Lemon Ave., Monrovia, CA 91016. To whom correspondence may be addressed: Alex Nisthal, Xencor Inc., 111 W. Lemon Ave., Monrovia, CA 91016; or Stephen L. Mayo, Division of Biology and Biological Engineering, California Institute of Technology, Pasadena, CA 91125.

## Abstract

The accurate prediction of protein stability upon sequence mutation is an important but unsolved challenge in protein engineering. Large mutational datasets are required to train computational predictors, but traditional methods for collecting stability data are either low-throughput or measure protein stability indirectly. Here, we develop an automated method to generate thermodynamic stability data for nearly every single mutant in a small 56-residue protein. Analysis reveals that most single mutants have a neutral effect on stability, mutational sensitivity is largely governed by residue burial, and unexpectedly, hydrophobics are the best tolerated amino acid type. Correlating the output of various stability prediction algorithms against our data shows that nearly all perform better on boundary and surface positions than for those in the core, and are better at predicting large to small mutations than small to large ones. We show that the most stable variants in the single mutant landscape are better identified using combinations of two prediction algorithms, and that including more algorithms can provide diminishing returns. In most cases, poor in silico predictions were tied to compositional differences between the data being analyzed and the datasets used to train the algorithm. Finally, we find that strategies to extract stabilities from high-throughput fitness data such as deep mutational scanning are promising and that data produced by these methods may be applicable toward training future stability prediction tools.

**Significance Statement:** Using liquid-handling automation, we constructed and measured the thermodynamic stability of almost every single mutant of protein G (Gβ1), a small domain. This self-consistent dataset is the largest of its kind and offers unique opportunities on two fronts: (*i*) insight into protein domain properties such as positional sensitivity and incorporated amino acid tolerance, and (*ii*) service as a validation set for future efforts in protein stability prediction. As Gβ1 is a model system for protein folding and design, and its single mutant landscape has been measured by deep mutational scanning, we expect our dataset to serve as a reference for studies aimed at extracting stability information from fitness data or developing novel high-throughput stability assays.

---

Thermodynamic stability is a fundamental property of proteins that significantly influences protein structure, function, expression, and solubility. Efforts to identify the molecular determinants of protein stability and to engineer improvements have thus been crucial in the development and optimization of a wide range of biotechnology products, including industrial-grade enzymes, antibodies, and other protein-based therapeutics and reagents (1-3). The ability to reliably predict the effect of mutations on protein stability would greatly facilitate engineering efforts, and much research has been devoted to developing computational tools for this purpose (4-9). Understanding how mutations affect stability can also shed light on various biological processes, including disease and drug resistance (10). More than 100,000 genetic variants have been associated with human disease (11) thanks to recent advances in genotyping and next generation sequencing, demonstrating a large need for fast and accurate stability prediction.

However, the accurate prediction of the impact of an amino acid substitution on protein stability remains an unsolved challenge in protein engineering. Correlation studies have shown that computational techniques can capture general trends, but fail to precisely predict the magnitude of mutational effects (12, 13). The success of these techniques is dependent on the quality of the input structure, conformational sampling, the free energy function used to evaluate the mutant sequences, and importantly, the data used for training and testing (8, 12, 14). Traditionally, protein stability data are collected by generating and purifying a small set of selected protein variants for characterization via calorimetry or spectroscopically measured chemical or thermal denaturation experiments. Values typically determined include the chemical or thermal denaturation midpoint (*C*_m_ or *T*_m_, respectively), the free energy of unfolding (Δ*G*), and the change in Δ*G* relative to wild type (WT) (ΔΔ*G*). Although low-throughput, the widespread use of these methods has generated a wealth of protein stability data over time, which has shaped our current understanding of protein structure-function relationships (15-18). Much of this work has been aggregated in the ProTherm (19) database, commonly used as a training data resource. Until recently, ProTherm was the largest public source of thermodynamic protein stability data, containing over 25,000 entries from 1,902 scientific articles. The database has been critical to the development of a variety of computational tools, from knowledge-based potentials exclusively trained on experimental data (6) to physics-based potentials with atomic resolution (7) and everything in between. Unfortunately, the ProTherm website is no longer being supported. The ProTherm data are still available, however, in ProtaBank (20), a recently developed online database for protein engineering data (https://protabank.org).

Although training and validation datasets from ProTherm have been widely used, ProTherm data suffer from three flaws: (*i*) experimental conditions vary widely among entries, requiring manual filtering to obtain comparable data, which results in smaller datasets, (*ii*) little information is included on unfolded or alternatively folded sequences, precluding training on this type of mutational data, and (*iii*) results from alanine scanning mutagenesis are overrepresented, biasing the dataset toward large to small mutations. Thus, training or testing on ProTherm data may mask deficiencies in computational algorithms or result in predictions that are biased toward particular features of the dataset. As many of the stability prediction tools available today rely on experimental data from ProTherm, it is perhaps not surprising that none are very accurate and all perform about the same (12).

Comprehensive mutagenesis studies, with stabilities measured under fixed experimental conditions, could provide better training data. The low-throughput nature of traditional methods, however, makes the collection of stability data for large numbers of protein variants unfeasible. Several strategies have been devised to improve this process, including the use of genetic repressor systems (21), plate-based fluorescence assays (22, 23), differential scanning fluorimetry (24), and more recently, yeast-displayed proteolysis (25). Unfortunately, these approaches generally make compromises by either: (*i*) tying an easy-to-measure but indirect protein stability readout to large variant libraries, or (*ii*) addressing the throughput of stability determination, but not the laborious nature of variant generation and purification.

Here, we develop an automated method that addresses both of these issues and apply it to obtain thermodynamic stability data from the comprehensive mutagenesis of an entire protein domain—the 56-residue β1 domain of Streptococcal protein G (Gβ1). Gβ1 was chosen for its small size, high amount of secondary structure, and well-behaved WT sequence. Drawing both inspiration and methodology from structural genomics, we couple automated molecular biology procedures with a high-throughput plate-based stability determination method, resulting in a 20-fold increase in throughput over traditional bench-top methods. We applied our experimental pipeline to Gβ1 to produce a dataset that maintains constant experimental conditions, includes data on non-folded sequences, and features an unbiased mutational distribution over 935 unique variants covering nearly every single mutant of Gβ1. Data in hand, we examine positional sensitivity and amino acid tolerance, and evaluate several protein stability prediction algorithms and engineering strategies. Finally, we compare our dataset against one derived by deep mutational scanning (DMS), a technique that can generate large mutational datasets via functional selections and deep sequencing (26, 27), and explore whether stability data from DMS studies are applicable towards training future protein stability prediction tools.

## Results and Discussion

### Automated Site-Directed Mutagenesis and Stability Determination Pipeline Increases Throughput 20-Fold

Using laboratory automation, we constructed, expressed, and purified nearly every single mutant in Gβ1. The automated pipeline is illustrated in Fig. 1*A*. Each variant was constructed explicitly instead of by saturation mutagenesis so that mutants not found in the first pass could be more easily recovered. Variants were constructed using a megaprimer method that requires only one mutagenic oligonucleotide, thereby halving oligonucleotide costs. The thermodynamic stabilities of the generated variants were then determined using an improved version of our previously described plate-based chemical denaturation assay (22) (Fig. 1*B*). Enhancements include adaptation to automated liquid handling for increased speed and precision, and doubling the number of data points collected per curve to improve accuracy. Although the intent was to collect data on 19 amino acids at 56 positions for a total of 1,064 variants, a tradeoff was made in which mutations at the buried tryptophan (Trp) at position 43 (W43) were excluded to preserve the integrity of the Trp-based fluorescence assay. Also, mutations incorporating cysteine (Cys) or Trp were omitted to avoid oligomerization by disulfide formation and potential interference with W43, respectively. Thus, mutations to 17 of 19 possible amino acids were made at 55 of 56 positions, for a total of 935 single mutants.

**Fig. 1.**
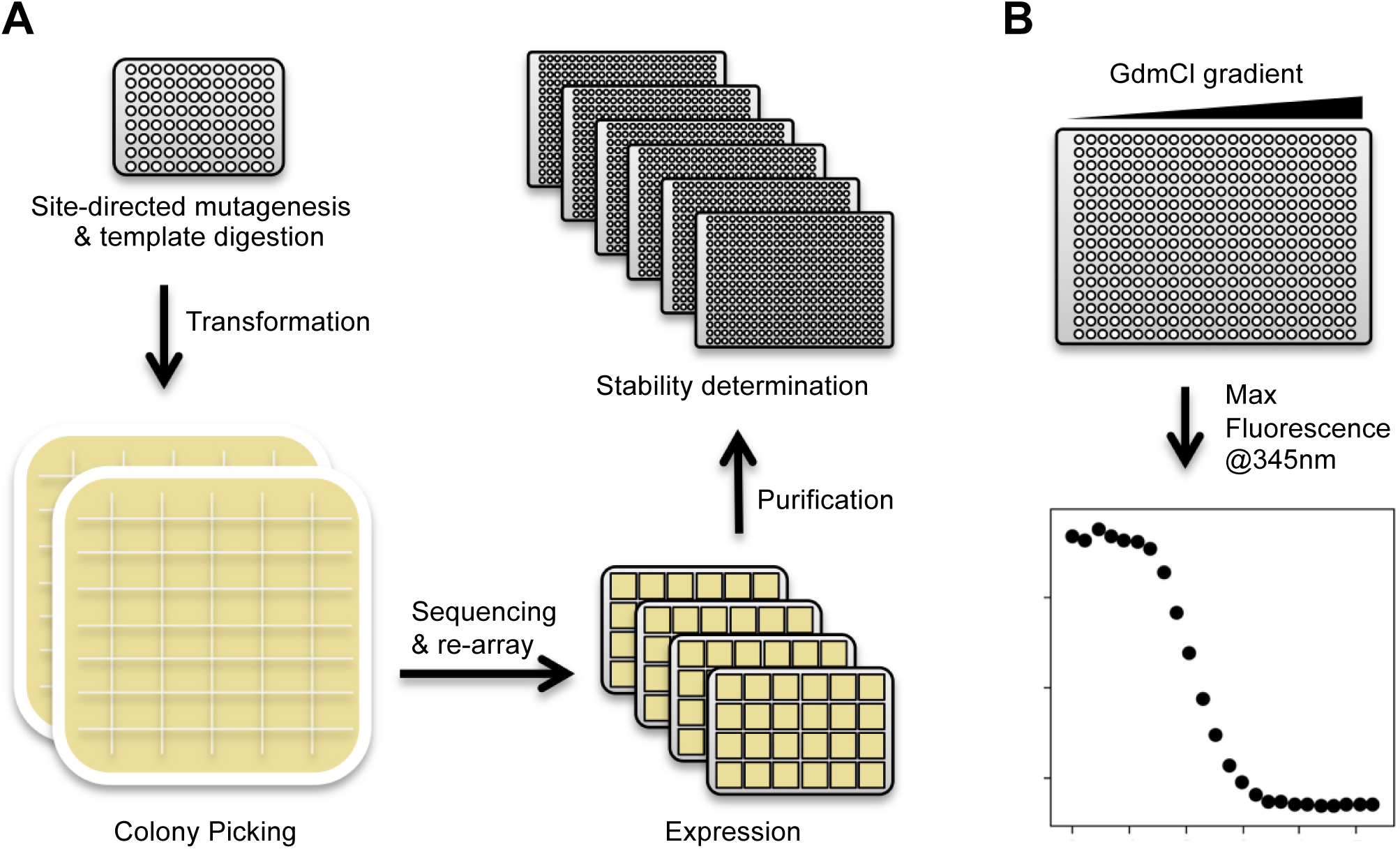
Automated site-directed mutagenesis and stability determination pipeline. (*A*) Modular protocols enable the rapid construction, sequence verification, and stability determination of single mutant variants. For illustrative purposes, each step shows the number of plates required for 96 individual reactions. Oligonucleotides direct mutagenesis reactions on 96-well PCR plates followed by bacterial transformation and plating onto 48-well agar trays. Individual colonies are picked, cultured, and re-arrayed after successful sequence validation. Confirmed variants are expressed in 24-well culture blocks and NiNTA purified. Each reaction was tracked via a database throughout the pipeline, allowing for method optimization. (*B*) Thermodynamic stability was determined by measuring Trp fluorescence in response to a 24-point GdmCl gradient. Each row of a 384-well plate is one protein stability curve from which the concentration of denaturant at the midpoint of the unfolding transition (*C*_m_) is directly measured. After estimating the slope of the curve (*m*-value), the change in the free energy of unfolding (ΔΔ*G*) of each variant relative to WT is calculated by taking the difference between the WT and mutant *C*_m_ values and multiplying by their mean *m*-value (see *SI Appendix*, Methods and *SI Appendix*, Fig. S2).

Each step of the workflow was developed as an independent module, allowing for optimization outside the full experimental pipeline. Modularization also permits flexible scheduling and parallelization, allowing modules to run multiple times per day. For comparison, eight days is a reasonable estimate for traditional procedures to construct, verify, express, purify, and measure the thermodynamic stability of 8 single mutants. Extrapolating to 935 variants (the number in this study), traditional procedures would take 935 days, or 2.5 years. In contrast, our platform can generate data on 935 variants in 5–6 weeks, a speedup of at least 20-fold.

### Stability Determination of Gβ1 Single Mutants

We measured the Trp fluorescence of each variant in response to a 24-point guanidinium chloride (GdmCl) gradient, thereby generating an unfolding curve (Fig. 1*B*) from which we determined the concentration of denaturant at the midpoint of the unfolding transition (*C*_m_) and the slope (*m*-value) (28). While ΔΔ*G* can be calculated in multiple ways (*SI Appendix,* Methods), a more precise method for our data takes the difference between the mutant and WT *C*_m_ values and multiplies it by their mean *m-*value 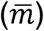 (29) as shown in the following equation:

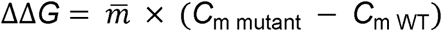

where the *m*-value was obtained with the linear extrapolation method (30). Using this equation, stabilizing mutations have positive ΔΔ*G* values, and destabilizing mutations have negative values. Of the 935 variants analyzed, 105 failed the assumptions of the linear extrapolation method (reversibility of folding/unfolding and two-state behavior) due to poor stability, presence of a folding intermediate, or no expression (*SI Appendix*, Fig. S1). The 830 variants that passed these criteria are referred to as the quantitative dataset, and the remaining 105 are referred to as the qualitative dataset. The single mutant stabilities (ΔΔ*G*s) for the entire dataset are shown as a heat map in Fig. 2.

**Fig. 2.**
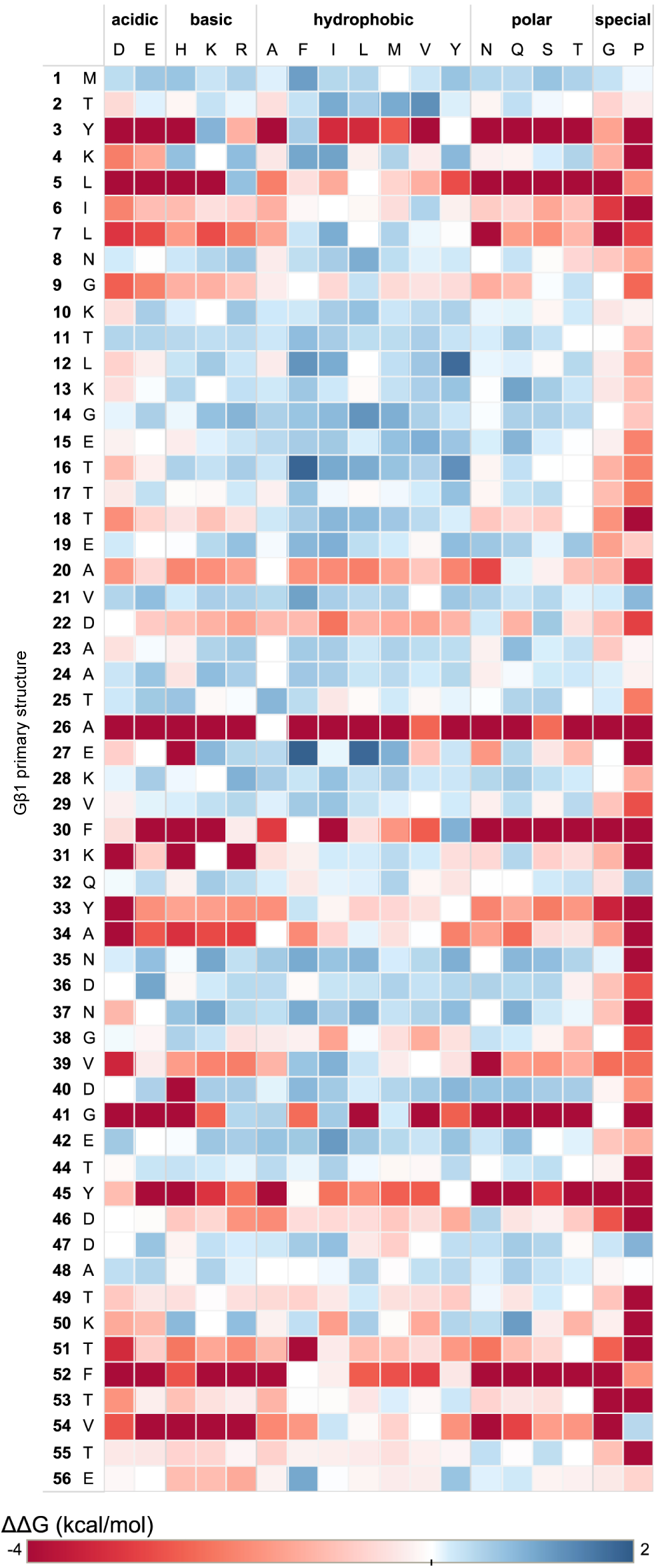
Single mutant thermodynamic stability landscape of Gβ1. The vertical axis of the mutational matrix depicts the primary structure of Gβ1, with the position and WT amino acid as columns. The horizontal axis depicts mutant amino acids examined in the study, grouped by amino acid type. Variants are colored by their determined ΔΔ*G* value where red is destabilizing, blue is stabilizing, and white is neutral. Self-identity mutations such as M1M have ΔΔ*G* = 0 and thus are colored white. Variants from the qualitative dataset are assigned an arbitrary value of –4 kcal/mol and are colored accordingly.

### Stability Distribution of Gβ1 Single Mutants Is Primarily Neutral

The ΔΔ*G* distribution of Gβ1 single mutants is primarily neutral (ΔΔ*G* of 0 ± 1 kcal/mol) with a long tail of destabilizing variants (Fig. 3*A*). The median of the quantitative dataset is 0.05 kcal/mol with an interquartile range of 1.0 kcal/mol (Fig. 3*C*), and the fraction of positive, neutral, and negative mutations is 3%, 68%, and 29%, respectively. If we assume the qualitative data contains only negative mutations, then our complete dataset shifts the fractions to 3%, 60%, and 37%, respectively. Summing the positive and neutral mutations, almost two thirds of the tested single mutants (63%) have at worst no effect on Gβ1 stability. The fraction of destabilizing mutations (37%) is on the low end compared to an experimental dataset of 1285 mutants from ProTherm, which shows that ∼50% of single mutants are destabilizing (ΔΔ*G* < 1 kcal/mol) (32, 33). The destabilizing fraction we obtained for Gβ1 would likely increase, however, upon making mutations to W43 and including Trp and Cys scanning variants as these residues are generally difficult to substitute in or out (34). Also, the Gβ1 domain itself may skew mutational outcomes as its small size results in a large surface-to-buried area ratio. This ratio likely contributes to fewer destabilizing mutations than larger proteins with larger cores, assuming that most core mutations are destabilizing (16, 21, 35, 36).

**Fig. 3.**
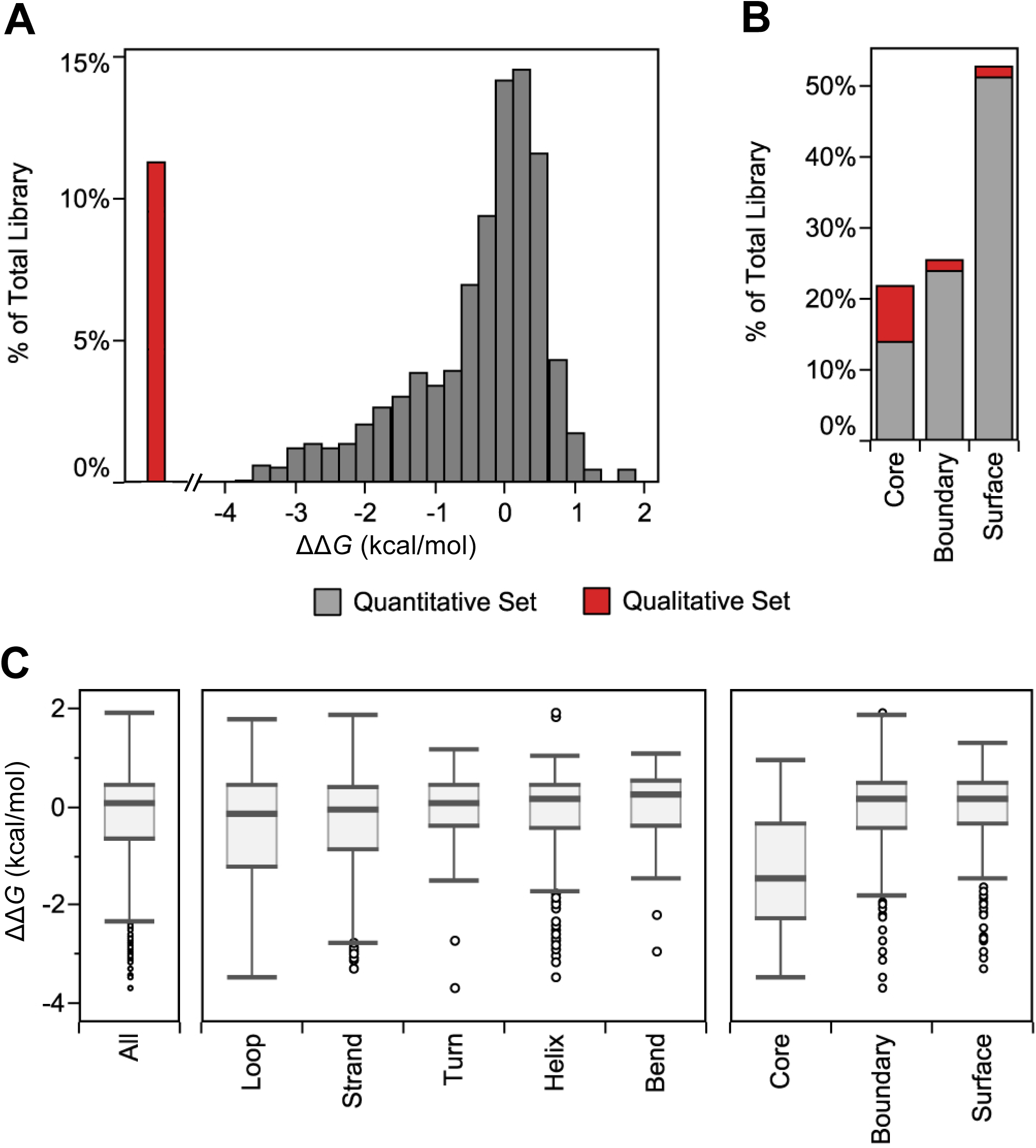
Stability distribution of Gβ1 single mutants. (*A* and *B*) The 935-member dataset is split between variants with quantitative data (gray) and those with only qualitative data (red) due to poor stability or misfolding. (*A*) The ΔΔ*G* distribution is split into 0.25 kcal/mol bins. Variants belonging to the qualitative dataset are shown to the left of the distribution, indicating ΔΔ*G*s < –4 kcal/mol. (*B*) Variants are binned into core, boundary, or surface using RESCLASS (4). (*C*) Box and whisker plots of the quantitative dataset describe the median, the quartile cutoffs, and the outlier cutoffs of the ΔΔ*G* distribution for all the residues (All), binned into secondary structure classifications as defined by DSSP (31), or binned by RESCLASS. Outliers are shown as unfilled circles and are defined as points that are 1.5 × interquartile range above or below the 3^rd^ quartile or 1^st^ quartile, respectively.

### Positional Sensitivity Is Governed by Residue Burial

The heat map in Fig. 2, which is organized by primary structure, allows for a granular look at the distribution of mutational stability. We observe two clear trends: (*i*) the mutational sensitivity (ΔΔ*G*) of the domain is largely determined by the position of the mutation, not the amino acid identity, unless (*ii*) the mutations are to glycine (Gly) or proline (Pro), for which most mutations are deleterious. Positions 3, 5, 26, 30, 41, 45, 52, and 54 are particularly sensitive to mutation. If we map the positional sensitivity (median ΔΔ*G* at each position) onto the Gβ1 structure (Fig. 4), we see that residues in the interior of the protein are more susceptible to destabilization. This is also observed when analyzing the distribution by tertiary structure, but not by secondary structure (Fig. 3*C*). That is, classifying residues into core, boundary, or surface with the RESCLASS algorithm (4) shows that the median ΔΔ*G* for core residues is ∼1.5 kcal/mol lower than that of the rest of the protein. In addition, the qualitative dataset, which contains mutants whose stabilities are difficult to measure or are fully unfolded, has 5-fold more core variants as compared to the boundary or surface, adding further support to this observation (Fig. 3*B*). Although this relationship has been observed with other datasets using a variety of proxies for protein stability (16, 21, 35, 36), this study provides a comprehensive analysis at the whole domain level with direct thermodynamic stability measurements.

**Fig. 4.**
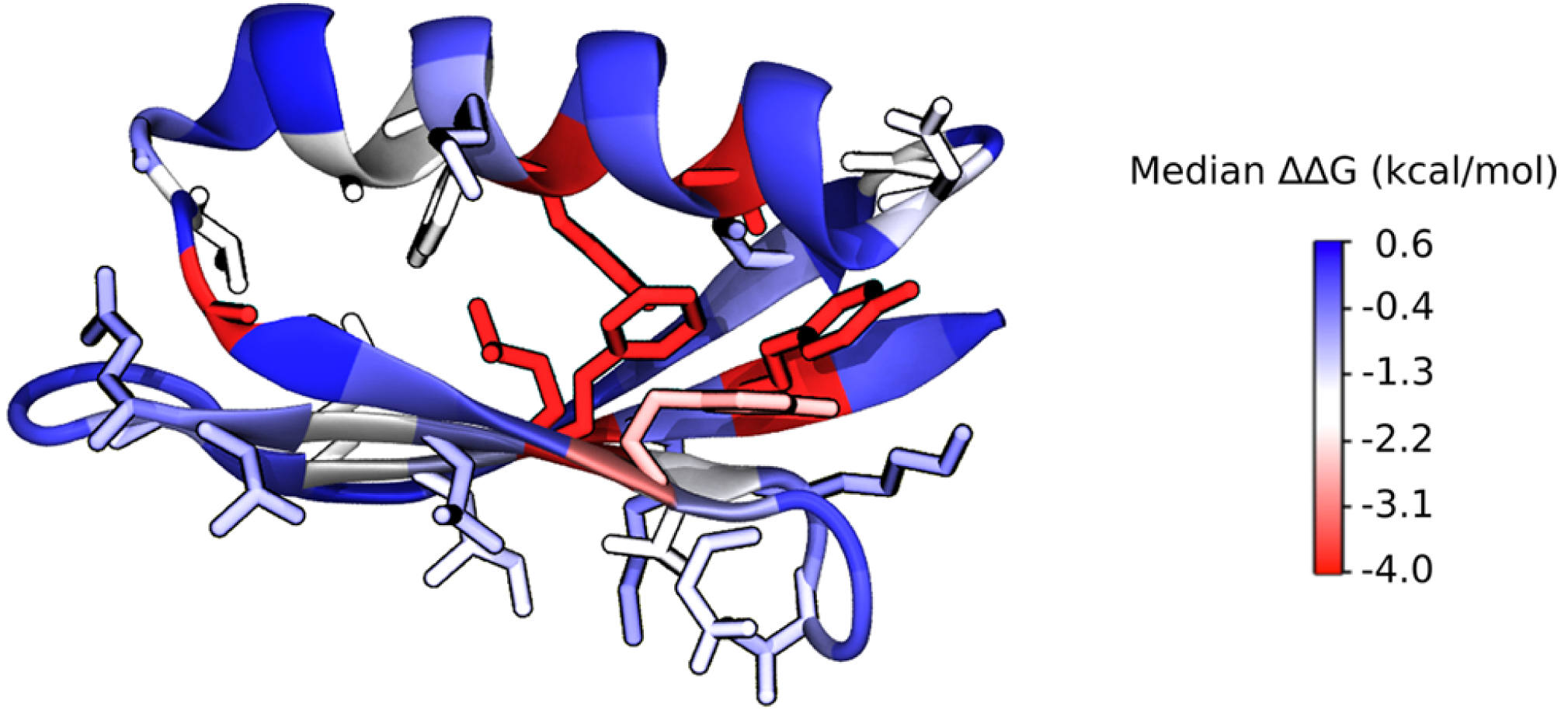
Positional sensitivity (median ΔΔ*G* at each position) of Gβ1. Gβ1 X-ray crystal structure (PDB ID: 1PGA) is colored by the positional sensitivity at each position. Sidechain atoms are shown for residues with a positional sensitivity score less than zero (destabilized).

As seen in Fig. 2, however, not all core positions behave the same, as some are more sensitive to mutation than others. For engineering purposes, it would be useful to identify specific protein attributes that could serve as quantitative predictors of positional sensitivity. We therefore performed linear regression with 10-fold cross validation on a large number of attributes that might impact protein stability. Attributes tested included measures of residue burial, secondary structure type/propensity, structural flexibility, and the change upon mutation of residue descriptors such as hydrophobicity, volume, and charge. The best individual predictors were measures of residue burial: depth of the Cβ atom (37, 38) and occluded surface packing (OSP)(39, 40), with correlation coefficients (*r*) of 0.82 and 0.76, respectively. This demonstrates that not all core positions are created equal, and that there is a direct relationship between how buried a position is and its sensitivity to mutation. Flexibility descriptors such as root mean squared fluctuations (RMSF) (from molecular dynamics simulations) or secondary structure descriptors such as *α*-helix propensity performed less well (*r* = 0.42 and 0.06, respectively). We repeated these analyses with sequence entropy (41) as an alternative metric of positional sensitivity, and the conclusions remain the same (Cβ depth and OSP were the two best predictors, with *r* = 0.81 and 0.78, respectively). Combinations of attributes were also tested, but these did not substantially improve predictability. Given the strong correlation between positional sensitivity and residue burial indicators like OSP and Cβ depth, calculation of these measures should be among the first tools employed when evaluating positions for substitution, provided structural information is available.

### Hydrophobics Are the Best Tolerated Amino Acid Type

A common practice in protein redesign and optimization is to restrict core residues to nonpolar amino acids and only allow polar amino acids at the surface. We tested the validity of this strategy with our quantitative dataset by calculating median ΔΔ*G* by incorporated amino acid and ranking the amino acids from worst tolerated to best tolerated across the entire domain (Fig. 5*A*). In general, the two worst amino acids for incorporation are Pro and Gly, which is unsurprising given their vastly different Ramachandran preferences compared to all other amino acids. Beyond secondary structure-breaking amino acids, the third worst tolerated amino acid, interestingly, is aspartic acid (Asp), which may be rationalized by the fact that it is very hydrophilic (42) and has one of the highest charge densities among the amino acids (43). Unexpectedly, hydrophobic amino acids, particularly isoleucine (Ile) and phenylalanine (Phe), are among the best tolerated residues across all Gβ1 positions. Even among surface positions, which make up over 50% of the dataset, Ile is the most favored individual residue, and hydrophobic amino acids as a whole are favored equally or better than the other amino acid types (Fig. 5*B*). The preference for hydrophobic amino acids extends to the chemically similar amino acid pairs, Asp/Glu and Asn/Gln, where the pair member containing the extra methylene is better tolerated across the domain (Fig. 5*A*) and in almost every RESCLASS environment (Fig. 5*C*). To determine if this observation is unique to Gβ1, we performed domain-wide in silico stability predictions (6, 44) on five compositionally diverse proteins, including Gβ1 (*SI Appendix*, Table S1). Remarkably, the calculations recapitulated our observations for Gβ1 and produced similar results for the other proteins, even across different RESCLASS types (*SI Appendix,* Fig. S3).

**Fig. 5.**
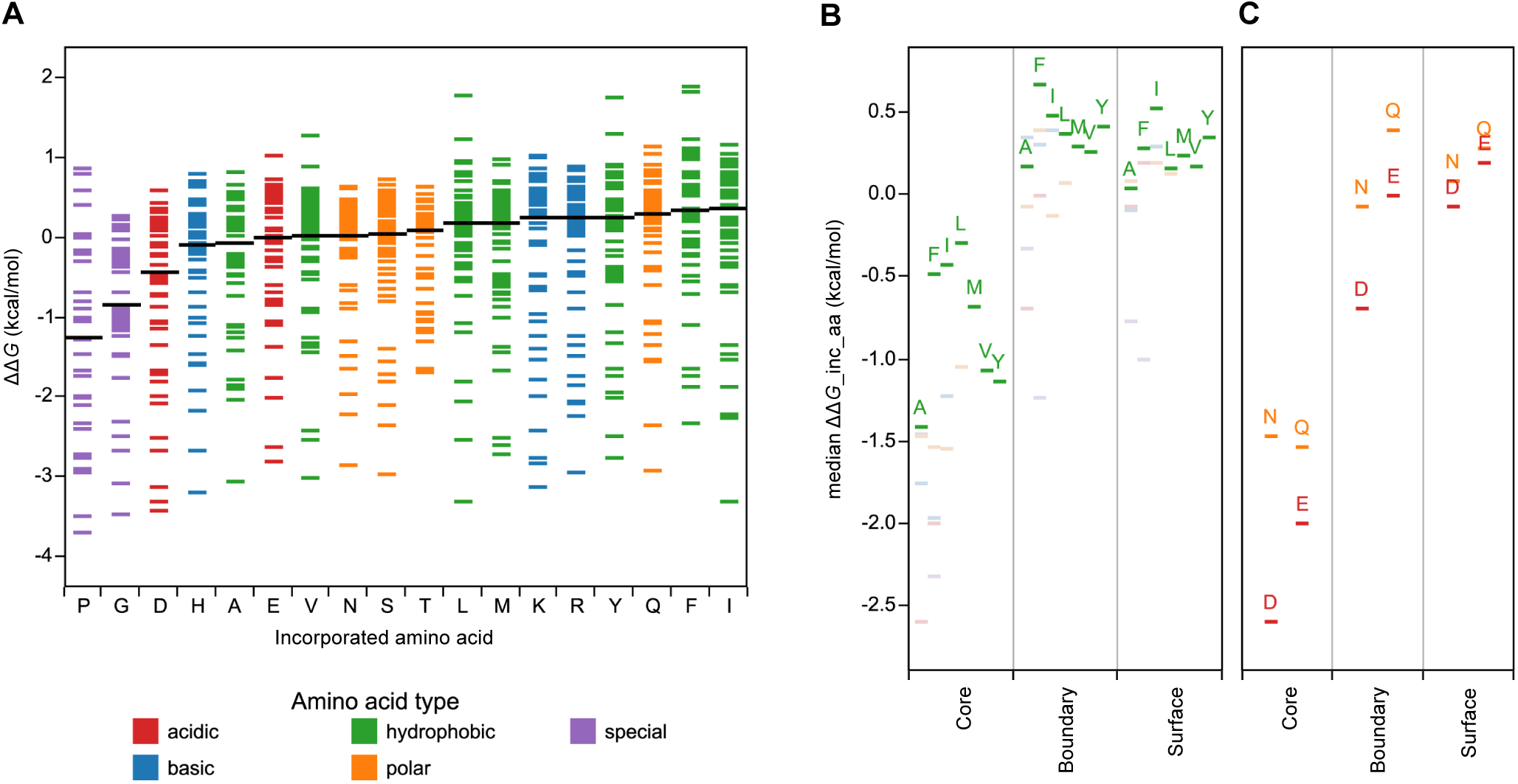
ΔΔ*G* distribution by incorporated amino acid. Amino acids are colored by physiochemical type. (*A*) Individual variants are shown as Gantt lines and distributed by the incorporated amino acid. The amino acid bins are ordered from left to right by the median ΔΔ*G* of each distribution (black lines). (*B*) Median ΔΔ*Gs* of the incorporated amino acid distribution grouped by RESCLASS (4). For clarity, only hydrophobic amino acids are labeled. (*C*) Median ΔΔ*Gs* of chemically similar pairs (D/E and N/Q), grouped by RESCLASS.

Several other experimental studies have also found that hydrophobic amino acids are well tolerated on the surface (45-49). The investigators attributed these findings to unique amino acid properties or structural contexts that enable these nonpolar mutations to stabilize the mutation site. However, our results suggest that non-position-specific increases in nonpolar surface area and volume are well tolerated, and the more the better. Larger hydrophobic amino acids like Ile, Phe, and Tyr are consistently ranked as the best tolerated, and smaller hydrophobics like Ala or Val do much worse across all three residue classes, including surface residues (Fig. 5*B*). Although multiple nonpolar mutations to the surface are still likely deleterious to protein stability and solubility (46), single mutations to hydrophobic amino acids should not be categorically excluded for stability optimization.

### Benchmarking Protein Stability Prediction Algorithms

We evaluated the ability of three stability prediction algorithms, PoPMuSiC (44), FoldX (7), and Rosetta (5, 8), to recapitulate the 830 ΔΔ*G* values in our quantitative dataset. To better understand the effect of training data on each algorithm’s performance, we compare the mutational composition of ΔΔ*G* datasets used in the development of each algorithm (Table 1). PoPMuSiC is a simplified-representation statistical energy function trained on a very large experimental dataset from ProTherm. FoldX is similarly trained, albeit with a smaller and more Ala biased dataset, and mixes all-atom physical potentials with weighted statistical terms. Rosetta also mixes statistical and all-atom physical potentials, but is trained to recover native sequence compositions for protein design. A recent study systematically explored the effect of 19 different Rosetta parameter sets on single mutant stability prediction (8), four of which are evaluated here. Three of the tested parameter sets use identical weights and terms but allow increasing amounts of backbone flexibility. That is, after sidechain repacking, the structure either undergoes no energy minimization, constrained backbone minimization, or unconstrained backbone minimization. Initially described as row 3, row 16, and row 19 (8), we refer to these parameter sets here as NoMin, SomeMin, and FullMin, respectively. The fourth Rosetta parameter set evaluated here (SomeMin_ddg) combines constrained minimization with optimized amino acid reference energies trained on single mutant ΔΔ*G* data from ProTherm, similar to FoldX and PoPMuSiC. Pearson correlation coefficients were used to evaluate algorithm performance overall (all mutations) as well as performance based on tertiary structure (RESCLASS) and volume change (Table 2). As energies from physical potentials can be dramatically skewed by atomic clashes, we excluded mutations with exceptionally high clash energies (clash outliers).

The Rosetta SomeMin method is the best performing algorithm overall with a Pearson correlation coefficient of 0.64 (Table 2). The other methods perform less well at *r* = 0.56 (PoPMuSiC) and *r* = 0.51 (FoldX). All of the algorithms scored lower on our dataset than previously reported on independent test sets, where *r* values of 0.69 (Rosetta SomeMin) (8), 0.67 (PopMuSiC) (44), and 0.64 (FoldX) (7) were obtained. Comparing the different Rosetta methods, we observe that increasing backbone flexibility decreases the number of clash outliers, but does not necessarily improve overall performance. The constrained minimization in SomeMin considerably improves the correlation over NoMin, but unconstrained minimization in FullMin shows diminishing returns in allowing increased flexibility, as observed previously (8). Significantly, the Rosetta SomeMin_ddg method performed worse than the SomeMin method (*r* = 0.54 and 0.64, respectively), demonstrating a limitation of training all-atom potentials with small, biased experimental datasets (Table 1).

If we look at the Pearson correlation coefficient by residue class, we find a general performance trend of boundary > surface > core. Except for Rosetta NoMin, which performs poorly across all categories, the all-atom algorithms exhibit very strong correlations in the boundary (*r* ≈ 0.7), with weaker correlations on the surface (*r* ≈ 0.5). In contrast, PoPMuSiC performs similarly across these two residue classes (*r* = 0.56 and *r* = 0.51, respectively). All algorithms do a poor job at predicting core mutations (*r* values range from 0.13 to 0.37), possibly because these mutations are more likely to lead to structural rearrangements that are not well captured by the algorithms (6-8). The significant differences in correlation accuracy observed here likely do not stem from deficiencies in training data, as the composition by residue class is fairly uniform across algorithms (Table 1).

The data were also analyzed by mutations that either reduce side chain volume (large to small, –VolΔ) or increase side chain volume (small to large, +VolΔ). Overall, across all methods, large to small mutations are better predicted than the inverse, which correlates with the composition of the training sets used in algorithm development (Table 1).

All algorithms were also evaluated by the Spearman correlation coefficient to minimize penalties on skewed energies and instead reward correct rank ordering. The differences found with the Pearson method on the overall dataset are no longer observed (*SI Appendix*, Table S2). PoPMuSiC and all the Rosetta methods perform about the same, with FoldX performing less well. However, the performance trend between residue classes is retained with boundary > surface > core, and the performance edge for large to small mutations is widened when evaluated by the Spearman coefficient. Because mutations that remove substantial volume often create a destabilizing cavity (50), the direction of the stability change of large to small mutations is more easily predicted and indeed captured by all of the algorithms equally well. The small to large mutation type can have very different outcomes (stabilized backbone accommodation or under/over-packed destabilization) and thus is harder to rank, much less predict accurately, as observed here. The trend in the volume change data subset demonstrates why stability predictors often feature favorable correlation coefficients on their test sets, which nearly always contain a bias towards mutations to small amino acids like Ala, as observed in Table 1.

### Practical Stability Engineering with in Silico Methods

As shown above and previously (12), highly accurate stability prediction (*r* > 0.8) is beyond current algorithms. However, this limitation has not prevented the successful application of in silico tools to stabilize proteins and engineer protein interaction specificity (51). One common approach is to: (*i*) generate stability predictions for every single mutant of a domain, (*ii*) filter the stability predictions by an arbitrary ΔΔ*G*(predicted) cutoff, (*iii*) experimentally verify the small number of mutants above the cutoff, and (*iv*) combine the hits. Here, our objective is to identify the in silico method that best performs this task on our Gβ1 dataset. That is, determine which algorithm recovers the greater number of stable variants (i.e., hits) near the top of its own predicted single-mutant list. We do this by calculating a couple of metrics across the two sorted lists of experimental and predicted variants, and, starting from the most stable variant, sequentially increase the number of mutants (*N*) that are compared. The first metric, % enrichment (%*E*), records the percent overlap between a list of experimentally verified mutants and a list of in silico predictions:

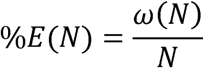

where *ω* (*N*) is the number of mutants found in both the experimental and predicted lists when *N* mutants are compared. The second metric, positive predictive value (*PPV*), first classifies the experimental dataset into “good” variants with ΔΔ*G* > 0 and “bad” variants with ΔΔ*G* ≤ 0, and then uses receiver operating characteristic (ROC) methods to calculate the fraction of true positives out of all positive predictions,

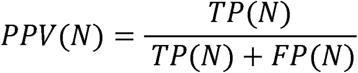

where *TP(N)* is the number of true positives when comparing lists of *N* mutations and *FP(N)* is the number of false positives when comparing lists of *N* mutations. Although both methods focus on positive predictions, %*E* is more sensitive to how stability algorithms order their comprehensive single mutant predictions, whereas *PPV* will give a favorable score as long as the mutants predicted are classified as “good” (ΔΔ*G* > 0).

Values of %*E* and *PPV* as a function of the number of mutants (*N*) in the comparison, or %*E*(*N*) and *PPV*(*N*), were calculated for FoldX, PoPMuSiC, and the four Rosetta methods. All combinations of the in silico methods were also tested by taking the mean of the predictions and then calculating %*E*(*N*) and *PPV*(*N*) as before. Focusing on the top 175 variants, which correspond to ΔΔ*G* > 0.5, we observe that the output values in general improve as the number of compared variants is increased (Fig. 6 and *SI Appendix*, Fig. S4). This result is expected, as testing more variants will increase the chances of identifying useful mutations. Both metrics indicate that Rosetta NoMin as a single algorithm returns the highest number of stabilizing mutations over the majority of variant cutoffs. Upon limiting *N* to the top 20 variants, the Rosetta methods with backbone flexibility outperform Rosetta NoMin. This result advocates for doing more computational modeling when experimental bandwidth is limiting. When considering combinations of two algorithms over the top 175 variants, the best performers are PoPMuSiC with any Rosetta protocol except SomeMin_ddg. These combinations have a higher %*E*(*N*) and *PPV*(*N*) than any single or combination of two algorithms when measuring the area under the curve (AUC) (*SI Appendix*, Fig. S5). Related work corroborates this result, showing that combinations of FoldX with other algorithms outperform FoldX alone in predicting the best single mutants (46, 51). However, no three-algorithm combinations significantly outperform the best two-algorithm combinations, indicating diminishing returns upon adding more algorithms, or at least the algorithms tested here.

**Fig. 6.**
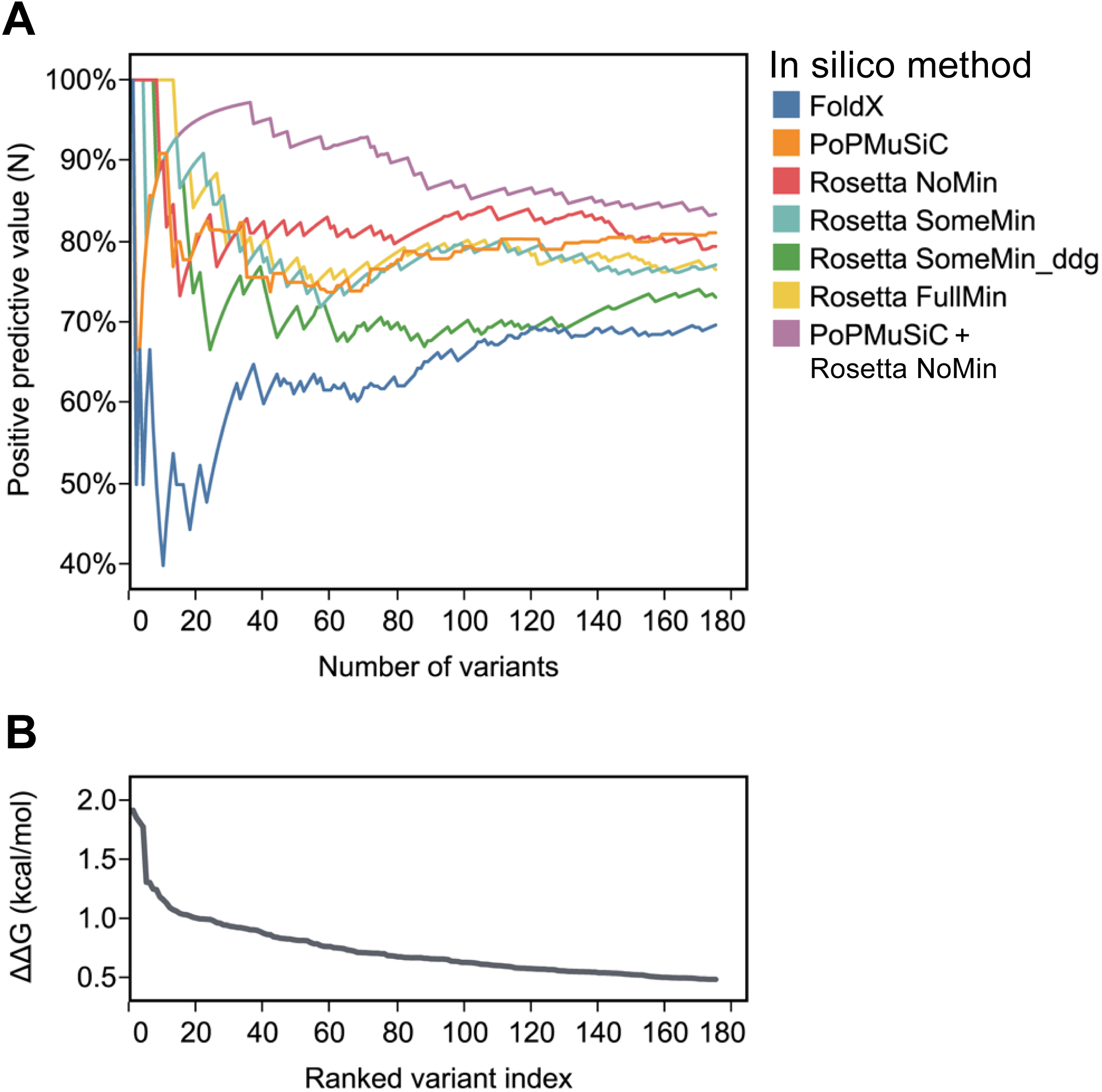
Comparing stability prediction algorithms by positive predictive value, *PPV*. (*A*) *PPV*(*N*), as defined in the text, is plotted as a function of the number of variants included in the list comparison. Only the top 175 Gβ1 single mutants are shown, sorted by ΔΔ*G*. Each of the single algorithms and, for simplicity, only the best two-algorithm combination (PoPMuSiC+Rosetta NoMin) are shown, colored according to the legend. (*B*) As a reference, experimental ΔΔ*G* values are plotted as a function of the ranked variant index, a sorted list of the stability distribution.

When benchmarked against our unbiased experimental data, FoldX alone performed poorly regardless of the value of the ΔΔ*G* cutoff, but especially in the top 20 predicted variants (Fig. 6 and *SI Appendix*, Fig. S4). Combining FoldX with Rosetta NoMin showed no improvement in AUC over Rosetta NoMin alone (*SI Appendix*, Fig. S5), demonstrating that combinations are sensitive to the quality of the input algorithms. Thus, it is perhaps not surprising that any three-algorithm combination involving FoldX failed to improve performance over the two-algorithm combinations. Similarly, the Rosetta SomeMin_ddg method lagged behind the other Rosetta-based methods as a single algorithm and in all higher combinations (*SI Appendix*, Fig. S5). As FoldX and Rosetta SomeMin_ddg are both all-atom potentials trained on single mutant data, we surmised that their respective training data sets were influencing their performance, as also shown by the correlative metrics. Indeed, the top 175 variants of the Gβ1 single mutant landscape are enriched 3 to 1 in small-to-large mutations, a mutation class that is vastly underrepresented in the training sets for FoldX and Rosetta SomeMin_ddg (Table 1).

### Comparing with Deep Mutational Scanning Studies

By coupling high-throughput functional selections with next generation sequencing, DMS can provide mutational data on thousands or even millions of variants with relatively little experimental effort (26, 27). This technology is being applied to an increasing number of proteins and has the potential to supply a wealth of new data to train stability prediction tools(52), provided the DMS technique can be properly validated. Serendipitously, a DMS study was performed on every single mutant and nearly every double-mutant of Gβ1 (53), allowing for a direct comparison with the thermodynamic stability data presented here. Using a selection based on binding to IgG Fc, Olson et al. found that fitness values obtained using binding affinity enrichment (ln *W*) correlated very poorly (*r* = 0.013) with ΔΔ*G* values reported in the literature for 82 single mutants (ΔΔ*G*_lit). When we compared ln *W* with the ΔΔ*G* values from our larger set of 830 single mutants, we found a better, but still small correlation (*r* = 0.19) (*SI Appendix*, Fig. S6*A*).

To address this issue, Olson et al. devised a strategy to estimate single mutant stabilities from their DMS fitness data. This approach requires identifying destabilized mutational backgrounds using double mutant fitness data so that the functional effect of a second mutation in these backgrounds could be used to compute single mutant ΔΔ*G*s. They identified five background mutations that produced a large correlation (*r* = 0.91) with ΔΔ*G*_lit and later demonstrated an approach [see Wu et al. (54)] that avoids the need for pre-existing stability data. In Fig. 7*A*, we plot our experimental ΔΔ*G*s vs. those predicted using the Wu et al. method (ΔΔ*G*_Wu) for 794 single mutants. The correlation (*r* = 0.60) is significantly lower than the value obtained using the smaller ΔΔ*G*_lit dataset (*r* = 0.91). A closer look at the 82 mutants in ΔΔ*G*_lit reveals a relatively small % of mutations in the core and a bias towards Ala substitutions, resulting in a dataset that does not reflect the breadth of possible mutations in the entire domain (Table 1). As seen in *SI Appendix*, Fig. S6*B*, the limited number of mutants in ΔΔ*G*_lit masks the lower correlation between ΔΔ*G* and ΔΔ*G*_Wu by serendipitously avoiding off-diagonal single mutants.

**Fig. 7.**
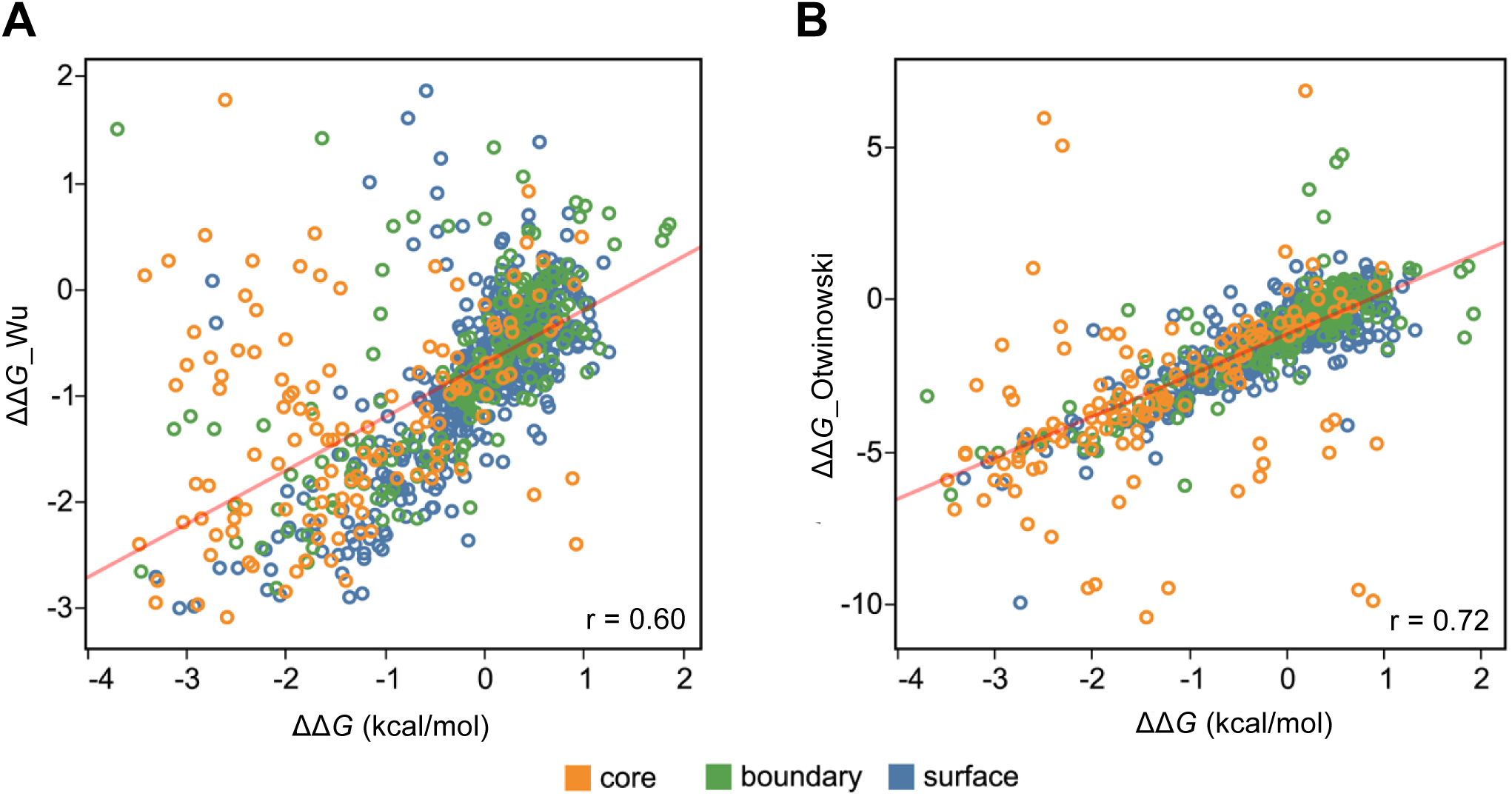
Comparing experimental ΔΔ*G*s with predictions obtained from DMS fitness data. Gβ1 single mutant stabilities from our experimental quantitative dataset (ΔΔ*G*) are plotted against (*A*) ΔΔ*G* values predicted from the DMS data using the Wu et al. method (54) (ΔΔ*G_*Wu) (*n* = 794) or (*B*) *E*_folding values predicted from the DMS data using Otwinowski’s three-state thermodynamic model (55) (ΔΔ*G_*Otwinowski) (*n* = 812). In both cases, DMS data is from Olson et al. (53). Points are colored by RESCLASS (4) values. A linear regression line is shown in red, and the correlation coefficient is shown in the lower right of each plot.

A recent report by Otwinowski reanalyzed the Olson et al. fitness data with a method based on a thermodynamic model describing three states (bound-folded, unbound-folded, and unfolded) that avoids the need for preexisting mutational or structural data (55). The method calculates distinct energies for folding (*E*_folding) and binding (*E*_binding). We compare the *E*_folding energy (ΔΔ*G*_Otwinowski) with our experimental ΔΔ*G* values in Fig. 7*B*, which shows an improved correlation (*r* = 0.72) over the Wu et al. method (*r* = 0.60). In *SI Appendix*, Table S3 analyzes the correlations for the two methods by residue class, volume change, and polarity change. The ΔΔ*G*_Otwinowski energy yields better correlations across the board, with the core continuing to show a significantly lower correlation. Thus, although DMS fitness data are poorly correlated with thermodynamic stability, simple biophysical models can be constructed that lead to significantly improved correlations. We expect that large, comprehensive datasets containing thermodynamic measurements such as those provided here will facilitate the development of improved methods to extract biophysical quantities (e.g., stability and binding) from fitness data, thus greatly expanding the utility of DMS and other deep sequencing techniques.

## Discussion

We described an automated chemical denaturation methodology that produces high quality thermodynamic stability data at a throughput that enables the near total site-saturation mutagenesis of small protein domains. Although other low-cost methods such as thermal challenge assays or differential scanning fluorimetry can also provide useful data, and deep sequencing approaches such as DMS can streamline the entire process, these methods do not directly report thermodynamic information. The automated pipeline described here makes gathering accurate thermodynamic stability data at a large scale feasible. The broad, unbiased nature of our near complete Gβ1 single mutant study provides an important dataset for examining domain-wide trends, evaluating stability prediction tools, and validating methods to extract stability values from DMS results. In addition, our analysis highlights the impact that training sets can have on computational predictors of stability.

We found that while the stability distribution of our Gβ1 dataset features a long tail of destabilizing variants, most mutations (68%) are neutral. However, if variants without quantitative data and those omitted for technical reasons are assigned negative outcomes, destabilizing variants make up 45% of the 1,064 possible single mutants of Gβ1, approaching predicted published values (32). Other trends followed conventions, with mutations to Gly, Pro, and core positions almost always being deleterious. However, not all core positions show the same degree of sensitivity, as measures of residue burial such as Cβ atom depth and OSP were found to best correlate with median ΔΔ*G* at each position. Although the correlation of residue burial with individual ΔΔ*G* measurements was previously reported for a collection of variants across many proteins (37), our domain-wide dataset allows the position-specific nature of the relationship to be fully observed. Similarly, using our unique dataset to calculate median ΔΔ*G* by incorporated amino acid reveals an unexpected tolerance for large hydrophobic amino acids. This preference extended across tertiary structure, and stability predictions on four other proteins confirmed this trend.

Evaluating three stability prediction algorithms against our dataset, we found that all performed moderately and recapitulated the general trends of the data. The flexible backbone Rosetta method (SomeMin) provided the best overall Pearson correlation (*r* = 0.64), but all of the Rosetta methods and PoPMuSiC performed equally by the Spearman rank correlation coefficient. Except for PoPMuSiC, all methods showed higher correlations in the boundary than on the surface (*r* = ∼0.7 and ∼0.5, respectively), all showed consistently lower correlations for mutations at core positions (*r* = 0.13 to 0.37), and nearly all were better at predicting large to small mutations than small to large ones.

Overall, the Rosetta SomeMin method was the most accurate stability algorithm tested here. It gives the best Pearson correlation for nearly every mutational category and is near the top in non-parametric methods as well. However, Rosetta SomeMin, and to a greater extent, Rosetta FullMin, require the most computational resources. For lower computational cost, PoPMuSiC provides the next best correlation coefficients on our quantitative dataset. For identification of the most stable single mutants, the combination of PoPMuSiC and Rosetta NoMin gave the best overall performance, and their excellent individual computational efficiencies should only increase their popularity. The combination was at or near the top in each of the metrics tested, yielding enrichment values over 30% and *PPV*s over 90% after analyzing the predicted top 175 variants.

DMS holds great promise as an extremely high-throughput method for obtaining mutational data for entire protein domains. However, correlating the fitness data to thermodynamic quantities such as stability is not straightforward, given that the selection method provides only an indirect measure of stability. In comparing our ΔΔ*G* values against strategies designed to extract stabilities from high-throughput fitness data, we find that a simple thermodynamic model that distinguishes binding and folding energies results in a Pearson correlation coefficient of *r* = 0.7. Even higher correlations are achieved by omitting core variants, and this strategy could yield useful training sets in the near term.

Beyond the engineering insights described here, it is our hope that our single mutant dataset of thermodynamic stabilities will prove to be a powerful validation set for use in developing better stability prediction tools and better methods for deriving stabilities from high-throughput fitness data.

## Materials and Methods

### Liquid Handling Robotics

A 2-meter Freedom EVO (Tecan) liquid-handling robot was used to automate the majority of the experimental pipeline. The instrument includes an eight-channel fixed-tip liquid-handling arm, a 96 disposable-tip single-channel liquid-handling arm, and a robotic plate-gripping arm. The robot’s deck features a fast-wash module, a refrigerated microplate carrier, a microplate orbital shaker, a SPE vacuum system, an integrated PTC-200 PCR machine (Bio-Rad Laboratories), stacks and hotels for microplates, and an integrated Infinite M1000 fluorescence microplate reader (Tecan). All molecular biology methods were developed de novo and optimized as necessary.

### Site-Directed Mutagenesis

The Gβ1 gene, with an N-terminal hexahistidine tag, was inserted into pET11a under control of an IPTG inducible T7 promoter. Mutagenic oligonucleotides were ordered from Integrated DNA Technologies in a 96-well format (150 µM concentration, 25 nmole scale) and purified by standard desalting. The site-directed mutagenesis reaction was performed in two parts: (*i*) diluted mutagenic oligonucleotides were mixed with a master mix solution composed of Hot-start Phusion DNA polymerase (NEB), GC Phusion buffer, dNTPs, the plasmid template, and the non-mutagenic flanking oligonucleotide, followed by (*ii*) mixing ¼ of the first step product with a similar master mix solution that omits the flanking oligonucleotide. The PCR cycling conditions for the two parts were: (*i*) a 30 s preincubation at 98 °C followed by 15 thermocycling steps (98 °C, 8 s; 62 °C, 15 s; 72 °C, 20 s), and (*ii*) a 30 s preincubation at 98 °C followed by 25 thermocycling steps (98 °C, 8 s; 72 °C, 3 min) followed by a final extension step at 72 °C for 5 min. After mutagenesis, samples were mixed into an 8%-by-volume Dpn1 (NEB) digestion reaction (37 °C, 1 h) to remove the parental template plasmid.

During development, reactions were diagnosed by E-Gel 96 (Invitrogen) electrophoresis systems, with loading performed by the liquid-handling robot. Visualization of the desired first-step and second-step products would guarantee positive mutagenesis. Almost 85% of all site-directed mutagenesis reactions were successful in the first pass.

### Bacterial Manipulation and Sequence Verification

Dpn1 digested products were mixed with homemade chemically competent BL21 Gold DE3 cells (56) in a 20 µL total reaction volume, and incubated at 4 °C for 10 min. After heat shock (42 °C, 45 s) on the PCR machine, the bacterial transformations were recovered by adding 100 µL of SOC media, and shaken off robot at 1200 rpm for 1 h at 37 °C on a microplate shaker (Heidolph).

The transformation mixtures were plated by the liquid-handling robot onto 48-well LB agar Qtrays (Genetix) and spread by sterile beads (56). The Qtrays were incubated off robot for 14 h at 37 °C. For each mutagenesis reaction, eight colonies were picked by a colony-picking robot (Qbot, Genetix) into 384-well plates (Genetix) filled with LB/10% glycerol. The 384-well receiving plates were incubated overnight at 37 °C, after which 2 of the 8 cultures per mutagenesis reaction were used to inoculate 96-well microplates containing LB/10% glycerol. These 96-well glycerol stock plates were grown overnight at 37 °C, replicated, and sent to Beckman Genomics for sequencing.

After analyzing the sequencing data, missing library members could be recovered either by sending more picked colonies from the 384-well receiving plate, or by redoing the mutagenesis reaction with different PCR conditions. Once all of the mutants were constructed, work lists were generated for the liquid-handling robot to cherry-pick from the replicated 96-well glycerol stock plates and inoculate into 96-well master stock plates containing LB/10% glycerol. The master stock plates were then incubated overnight at 37 °C and frozen at –80 °C until needed.

### Protein Expression and Purification

Small volumes from replicated master stock plates were used to inoculate 5 mL of Instant TB auto-induction media (Novagen) in 24-well round-bottom plates (Whatman). The 24-well plates were incubated overnight, shaking at 250 rpm, at 37 °C. The expression cultures were then pelleted, lysed with a sodium phosphate lysis buffer solution (pH 8) containing CelLytic B (Sigma Aldrich), lysozyme, and HC benzonase (Sigma Aldrich). Lysates were then added directly to 96-well His-Select Ni-NTA resin filter plates (Sigma Aldrich) and processed off-robot by centrifugation. His-tagged protein was washed and eluted in sodium phosphate buffer (pH 8) containing 0 mM and 100 mM imidazole, respectively. Protein samples were diluted five-fold into sodium phosphate buffer (pH 6.5), thereby diluting the final amount of imidazole in each sample before stability determination.

### Plate-Based Stability Assay

Large volumes (0.2 L) of each concentration of a 24-point gradient of GdmCl in sodium phosphate buffer (pH 6.5) were constructed using graduated cylinders and dispensed into 96-well deep-well plates by a multi-channel pipettor. These reagent reservoirs, along with the liquid-handling robot, greatly simplified and sped up the stability assay previously described (22). Each stability assay was comprised of 24 individual solutions containing 1 part purified protein to 4 parts GdmCl/buffer solution, and measured by the integrated plate reader for Trp fluorescence (Ex: 295 nm, Em: 341 nm). The assay employed 384-well UV-Star plates (Greiner) that allowed 16 different protein mutants to be measured per plate, thus requiring 6 of these plates per 96-well master stock plate. All variants were measured 2–6 times. Data were analyzed as described previously (22).

### Positional Sensitivity

Positional sensitivity was evaluated via two metrics: (*i*) the median ΔΔ*G* value and (*ii*) sequence entropy. The median ΔΔ*G* value for each position was calculated by finding the median of ΔΔ*G* values for all mutations measured at *j*, where mutations in the qualitative dataset were assigned a ΔΔ*G* value of –4 kcal/mol. The sequence entropy at a position *j* was calculated as 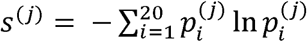 where 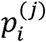, the probability of a given amino acid *i* at position *j* was determined by 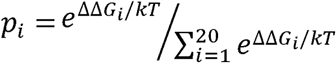. The WT residue was assigned a ΔΔ*G* value of zero and mutations in the qualitative dataset were assigned a value of –4 kcal/mol. The positional sensitivity at each position was visualized on the crystal structure of Gβ1 (PDB ID: 1PGA) using VMD (57).

### Protein Attributes

All structure-based attributes were calculated using the crystal structure of Gβ1 (PDB ID: 1PGA). Occluded surface packing (40) was calculated using software downloaded from http://pages.jh.edu/pfleming/sw/os/. Root mean square fluctuation (RMSF) was calculated over a 20 ns molecular dynamics trajectory in full solvent using NAMD (58). The depth of the Cβ atom was calculated by the RESCLASS algorithm (4) to decide core, boundary, and surface residues. Linear regression with 10-fold cross validation was performed with scikit-learn (59) to identify attributes that correlate highly with positional sensitivity. Recursive feature elimination was also performed with scikit-learn using a ridge estimator, and 5-fold cross validation was performed to evaluate combinations of attributes. Recursive feature elimination was also performed with scikit-learn to evaluate combinations of attributes.

### Stability Prediction Algorithms

The crystal structure of Gβ1 (PDB ID: 1PGA) was used as the input structure for all algorithms. The webserver for PoPMuSiC version 3, located at http://www.dezyme.com, was used to perform a “Systematic” command on the Gβ1 crystal structure. A copy of FoldX (version 3.0, beta 5) was retrieved from http://foldx.crg.es. The crystal structure was prepared by using the “RepairPDB” command to perform Asn, Gln, and His flips, alleviate small van der Waals clashes, and optimize WT rotamer packing. Every mutant in the dataset was constructed through the “BuildModel” command, and the difference in energy between the WT reference and the corresponding mutant was averaged over five trials. A copy of Rosetta (version 3.3) was retrieved from http://www.rosettacommons.org. The ddg_monomer application was used to generate single mutant stability data from the Gβ1 crystal structure. We followed the available online documentation in order to prepare all necessary input files. Option sets described in the documentation pertain to the various Rosetta iterations tested in this paper (NoMin: low-resolution protocol; SomeMin: high-resolution protocol; FullMin: high-resolution protocol with an empty distance restraints file).

### Statistical Visualization and Analysis

All plots were generated using the software au (Seattle, WA). Custom python scripts were developed to calculate the large number of thermodynamic stability curve fits. Correlation coefficients (Pearson’s and Spearman’s) were calculated either in Tableau or in the software package R (version 3.2.2). The ROCR package for R was used for classification and receiver operator characteristic analysis (60).

### Data Availability

The ΔΔ*G* distribution of Gβ1 single mutants generated during this work is publicly available in ProtaBank (https://protabank.org), a protein engineering data repository, under the ID gwoS2haU3. All of the other data that support the conclusions of the study are available from A.N. upon request.

## Acknowledgements

A.N. thanks Jost Vielmetter for advice and feedback on the automated platform. S.L.M. acknowledges grants from the National Security Science and Engineering Faculty Fellowship program and the Defense Advanced Research Projects Agency Protein Design Processes program.

## Author contributions

A.N. and S.L.M designed the research. A.N. developed the experimental approach, performed all experiments, and managed the data. A.N., C.Y.W., and M.L.A. analyzed the data and wrote the paper with input from S.L.M.

## References

1. Bommarius AS, Paye MF (2013) Stabilizing biocatalysts. Chem Soc Rev 42:6534–6565.

2. Goldenzweig A, Fleishman S (2018) Principles of protein stability and their application in computational design. Annu Rev Biochem 87:105–129.

3. Rouet R, Lowe D, Christ D (2014) Stability engineering of the human antibody repertoire. FEBS Lett 588:269–277.

4. Dahiyat BI, Mayo SL (1997) De novo protein design: fully automated sequence selection. Science 278:82–87.

5. Das R, Baker D (2008) Macromolecular modeling with Rosetta. Annu Rev Biochem 77:363–382.

6. Dehouck Y, et al. (2009) Fast and accurate predictions of protein stability changes upon mutations using statistical potentials and neural networks: PoPMuSiC-2.0. Bioinformatics 25:2537–2543.

7. Guerois R, Nielsen JE, Serrano L (2002) Predicting changes in the stability of proteins and protein complexes: a study of more than 1000 mutations. J Mol Biol 320:369–387.

8. Kellogg EH, Leaver-Fay A, Baker D (2011) Role of conformational sampling in computing mutation-induced changes in protein structure and stability. Proteins 79:830–838.

9. Malakauskas SM, Mayo SL (1998) Design, structure and stability of a hyperthermophilic protein variant. Nat Struct Biol 5:470–475.

10. Pires DEV, Chen J, Blundell TL, Ascher DB (2016) In silico functional dissection of saturation mutagenesis: Interpreting the relationship between phenotypes and changes in protein stability, interactions and activity. Sci Rep 6:19848.

11. Burgess DJ (2015) Disease genetics: network effects of disease mutations. Nat Rev Genet 16:317.

12. Potapov V, Cohen M, Schreiber G (2009) Assessing computational methods for predicting protein stability upon mutation: good on average but not in the details. Protein Eng Des Sel 22:553–560.

13. Khan S, Vihinen M (2010) Performance of protein stability predictors. Hum Mutat 31:675–684.

14. Davey JA, Chica RA (2015) Optimization of rotamers prior to template minimization improves stability predictions made by computational protein design. Protein Sci 24:545–560.

15. Alber T (1989) Mutational effects on protein stability. Annu Rev Biochem 58:765–798.

16. Bowie JU, Reidhaar-Olson JF, Lim WA, Sauer RT (1990) Deciphering the message in protein sequences: tolerance to amino acid substitutions. Science 247:1306–1310.

17. Matthews BW (1993) Structural and genetic analysis of protein stability. Annu Rev Biochem 62:139–160.

18. Fersht AR, Serrano L (1993) Principles of protein stability derived from protein engineering experiments. Curr Opin Struct Biol 3:75–83.

19. Kumar MDS, et al. (2006) ProTherm and ProNIT: thermodynamic databases for proteins and protein-nucleic acid interactions. Nucleic Acids Res 34:D204–206.

20. Wang CY, et al. (2018) ProtaBank: A repository for protein design and engineering data. Protein Sci 27:1113–1124.

21. Markiewicz P, Kleina LG, Cruz C, Ehret S, Miller JH (1994) Genetic studies of the lac repressor. XIV. Analysis of 4000 altered Escherichia coli lac repressors reveals essential and non-essential residues, as well as “spacers” which do not require a specific sequence. J Mol Biol 240:421–433.

22. Allen BD, Nisthal A, Mayo SL (2010) Experimental library screening demonstrates the successful application of computational protein design to large structural ensembles. Proc Natl Acad Sci USA 107:19838–19843.

23. Aucamp JP, Cosme AM, Lye GJ, Dalby PA (2005) High-throughput measurement of protein stability in microtiter plates. Biotechnol Bioeng 89:599–607.

24. Lavinder JJ, Hari SB, Sullivan BJ, Magliery TJ (2009) High-throughput thermal scanning: a general, rapid dye-binding thermal shift screen for protein engineering. J Am Chem Soc 131:3794–3795.

25. Rocklin GJ, et al. (2017) Global analysis of protein folding using massively parallel design, synthesis, and testing. Science 357:168–175.

26. Araya CL, Fowler DM (2011) Deep mutational scanning: assessing protein function on a massive scale. Trends Biotechnol 29:435–442.

27. Fowler DM, Fields S (2014) Deep mutational scanning: a new style of protein science. Nat Methods 11:801–807.

28. Myers JK, Pace CN, Scholtz JM (1995) Denaturant m values and heat capacity changes: relation to changes in accessible surface areas of protein unfolding. Protein Sci 4:2138–2148.

29. Pace CN (1986) Determination and analysis of urea and guanidine hydrochloride denaturation curves. Methods Enzymol 131:266–280.

30. Santoro MM, Bolen DW (1988) Unfolding free energy changes determined by the linear extrapolation method. 1. Unfolding of phenylmethanesulfonyl alpha-chymotrypsin using different denaturants. Biochemistry 27:8063–8068.

31. Kabsch W, Sander C (1983) Dictionary of protein secondary structure: pattern recognition of hydrogen-bonded and geometrical features. Biopolymers 22:2577–2637.

32. Tokuriki N, Stricher F, Schymkowitz J, Serrano L, Tawfik DS (2007) The stability effects of protein mutations appear to be universally distributed. J Mol Biol 369:1318–1332.

33. Tokuriki N, Tawfik DS (2009) Stability effects of mutations and protein evolvability. Curr Opin Struct Biol 19:596–604.

34. Henikoff S, Henikoff JG (1992) Amino acid substitution matrices from protein blocks. Proc Natl Acad Sci USA 89:10915–10919.

35. Huang W, Petrosino J, Hirsch M, Shenkin PS, Palzkill T (1996) Amino acid sequence determinants of beta-lactamase structure and activity. J Mol Biol 258:688–703.

36. Rennell D, Bouvier SE, Hardy LW, Poteete AR (1991) Systematic mutation of bacteriophage T4 lysozyme. J Mol Biol 222:67–88.

37. Chakravarty S, Varadarajan R (1999) Residue depth: a novel parameter for the analysis of protein structure and stability. Structure 7:723–732.

38. Tan KP, Nguyen TB, Patel S, Varadarajan R, Madhusudhan MS (2013) Depth: a web server to compute depth, cavity sizes, detect potential small-molecule ligand-binding cavities and predict the pKa of ionizable residues in proteins. Nucleic Acids Res 41:W314–321.

39. Pattabiraman N, Ward KB, Fleming PJ (1995) Occluded molecular surface: analysis of protein packing. J Mol Recognit 8:334–344.

40. Fleming PJ, Richards FM (2000) Protein packing: dependence on protein size, secondary structure and amino acid composition. J Mol Biol 299:487–498.

41. Valdar WSJ (2002) Scoring residue conservation. Proteins 48:227–241.

42. Kyte J, Doolittle RF (1982) A simple method for displaying the hydropathic character of a protein. J Mol Biol 157:105–132.

43. Dixon DA, Lipscomb WN (1976) Electronic structure and bonding of the amino acids containing first row atoms. J Biol Chem 251:5992–6000.

44. Dehouck Y, Kwasigroch JM, Gilis D, Rooman M (2011) PoPMuSiC 2.1: a web server for the estimation of protein stability changes upon mutation and sequence optimality. BMC Bioinformatics 12:151.

45. Ayuso-Tejedor S, Abián O, Sancho J (2011) Underexposed polar residues and protein stabilization. Protein Eng Des Sel 24:171–177.

46. Broom A, Jacobi Z, Trainor K, Meiering EM (2017) Computational tools help improve protein stability but with a solubility tradeoff. J Biol Chem 292:14349–14361.

47. Cordes MH, Sauer RT (1999) Tolerance of a protein to multiple polar-to-hydrophobic surface substitutions. Protein Sci 8:318–325.

48. Machius M, Declerck N, Huber R, Wiegand G (2003) Kinetic stabilization of Bacillus licheniformis alpha-amylase through introduction of hydrophobic residues at the surface. J Biol Chem 278:11546–11553.

49. Poso D, Sessions RB, Lorch M, Clarke AR (2000) Progressive stabilization of intermediate and transition states in protein folding reactions by introducing surface hydrophobic residues. J Biol Chem 275:35723–35726.

50. Baase WA, Liu L, Tronrud DE, Matthews BW (2010) Lessons from the lysozyme of phage T4. Protein Sci 19:631–641.

51. Buß O, Rudat J, Ochsenreither K (2018) FoldX as protein engineering tool: better than random based approaches? Comput Struct Biotechnol J 16:25–33.

52. Araya CL, Fowler DM (2011) Deep mutational scanning: assessing protein function on a massive scale. Trends Biotechnol 29:435–442.

53. Olson CA, Wu NC, Sun R (2014) A comprehensive biophysical description of pairwise epistasis throughout an entire protein domain. Curr Biol 24:2643–2651.

54. Wu NC, Olson CA, Sun R (2015) High-throughput identification of protein mutant stability computed from a double mutant fitness landscape. Protein Sci 25:530–539.

55. Otwinowski J (2018) Biophysical inference of epistasis and the effects of mutations on protein stability and function. arXiv: 1802.08744v2. Preprint, posted Mar 30 2018.

56. Klock HE, Lesley SA (2009) The Polymerase Incomplete Primer Extension (PIPE) method applied to high-throughput cloning and site-directed mutagenesis. Methods Mol Biol 498:91–103.

57. Humphrey W, Dalke A, Schulten K (1996) VMD: visual molecular dynamics. J Mol Graph 14:33–38.

58. Phillips JC, et al. (2005) Scalable molecular dynamics with NAMD. J Comput Chem 26:1781–1802.

59. Pedregosa F, Varoquaux G, Gramfort A, Michel V, Thirion B (2011) Scikit-learn: Machine learning in Python. JMLR 12:2825–2830.

60. Sing T, Sander O, Beerenwinkel N, Lengauer T (2005) ROCR: visualizing classifier performance in R. Bioinformatics 21:3940–3941.

