## Supplementary material for "Protein stability engineering insights revealed by domain-wide comprehensive mutagenesis": Table 1

**Table 1. Mutational composition of protein stability datasets**

| ΔΔ*G* dataset | Total # | % surface | % boundary | % core | % +VolΔ | % −VolΔ | % Ala |
| --- | --- | --- | --- | --- | --- | --- | --- |
| FoldX training set (7) | 339 | 32 | 27 | 36 | 3 | 97 | 61 |
| FoldX test set (7) | 625 | 32 | 30 | 35 | 5 | 95 | 54 |
| PoPMuSiC training set (6) | 2644 | 26 | 32 | 40 | 33 | 67 | 28 |
| PoPMuSiC test set (6) | 350 | 21 | 27 | 48 | 40 | 60 | 26 |
| Rosetta test set (8) | 1210 | 32 | 30 | 38 | 16 | 84 | 47 |
| ΔΔ*G_*lit (53) | 82 | 71 | 21 | 8 | 52 | 48 | 20 |
| This dataset | 935 | 53 | 25 | 22 | 56 | 44 | 5 |
| Top 175 variants of this dataset | 175 | 63 | 32 | 5 | 78 | 22 | 3 |

Residues were classified as core, boundary or surface with the RESCLASS (4) algorithm. Mutations with mislabeled or non-standard PDB data (< 5%) were omitted from residue classification. +VolΔ, small to large mutations; **−**VolΔ, large to small mutations.
