## Supplementary material for "Protein stability engineering insights revealed by domain-wide comprehensive mutagenesis": Table 2

**Table 2. Algorithm performance by Pearson correlation**

|  |  | |  | |  | Pearson correlation coefficient (*r*) | | | | | |
| --- | --- | --- | --- | --- | --- | --- | --- | --- | --- | --- | --- |
| Algorithm | | Backbone minimization* | | Clash outliers^†^ | | Overall | Surface^‡^ | Boundary^‡^ | Core^‡^ | +VolΔ | −VolΔ |
| PoPMuSiC | |  | | 0 | | 0.56 | 0.51 | 0.56 | 0.33 | 0.43 | 0.64 |
| FoldX | |  | | 17 | | 0.51 | 0.42 | 0.68 | 0.17 | 0.46 | 0.56 |
| Rosetta^§^ | |  | |  | |  |  |  |  |  |  |
| NoMin | | None | | 22 | | 0.33 | 0.29 | 0.26 | 0.13 | 0.38 | 0.28 |
| SomeMin | | Constrained | | 17 | | 0.64 | 0.53 | 0.73 | 0.37 | 0.56 | 0.66 |
| SomeMin_ddg^¶^ | | Constrained | | 6 | | 0.54 | 0.49 | 0.68 | 0.15 | 0.46 | 0.66 |
| FullMin | | Unconstrained | | 3 | | 0.60 | 0.52 | 0.69 | 0.24 | 0.48 | 0.69 |

Predicted ΔΔ*G*s from stability algorithms were compared to experimental ΔΔ*G*s for Gβ1 single mutants in the quantitative dataset. Mutations with exceptionally high clash energies (clash outliers) were excluded when calculating each algorithm’s correlation coefficient. +VolΔ, small to large mutations; **−**VolΔ, large to small mutations.

*Level of backbone minimization after repacking for Rosetta methods.

^†^Number of mutations with a calculated repulsive energy > 2 standard deviations above the mean.

**^‡^**Residues are classified as core, boundary, or surface using RESCLASS (4).

^§^Rosetta parameter sets NoMin, SomeMin, and FullMin were initially described as row 3, row 16, and row 19, respectively (8).

^¶^Combines constrained backbone minimization with optimized reference energies trained on ProTherm single mutant ΔΔ*G* data.
