## Supplementary Material for "Protein stability engineering insights revealed by domain-wide comprehensive mutagenesis"

This PDF file includes:

Supplementary text

Figs. S1 to S6

Tables S1 to S3

References for SI reference citations

**Supplementary Information Text**

**Methods**

**ΔΔ*G* calculation.** The ΔΔ*G* of the 830 mutants in the quantitative dataset can be calculated in two ways: (*i*) by taking the difference between WT and mutant fitted Δ*G*(H_2_O) values given by the linear extrapolation method (1) (ΔΔ*G*(H_2_O)), or (*ii*) by taking the difference between WT and mutant *C*_m_ values and multiplying by their mean *m‑*value(2) (ΔΔ*G*(*m-*avg)). The ΔΔ*G*(*m-*avg) value avoids fitting issues near low denaturant values when estimating Δ*G*(H_2_O), but loses validity when variants greatly affect the *m*-value or the stability of the mutant protein (2, 3). As roughly 80% of the dataset has values ±1 kcal/mol away from WT, and the WT *m*-value is within 1 standard deviation of the mean *m*-value for the dataset, the *m*-avg method appears valid. After averaging repeat measurements, the two methods correlate extremely well (*r* = 0.99) with over 90% of the dataset differing by no more than 0.2 kcal/mol and none more than 1 kcal/mol (Fig. S2*A*). The only major difference between the methods is that the ΔΔ*G*(H_2_O) method is less precise than the ΔΔ*G*(*m-*avg) method (Fig. S2*B*), likely due to the uncertainties inherent in estimating Δ*G*(H_2_O) values. Thus, the mean of the ΔΔG(*m-*avg) values recorded for each variant (referred to as ΔΔ*G*) were used for all further analysis.

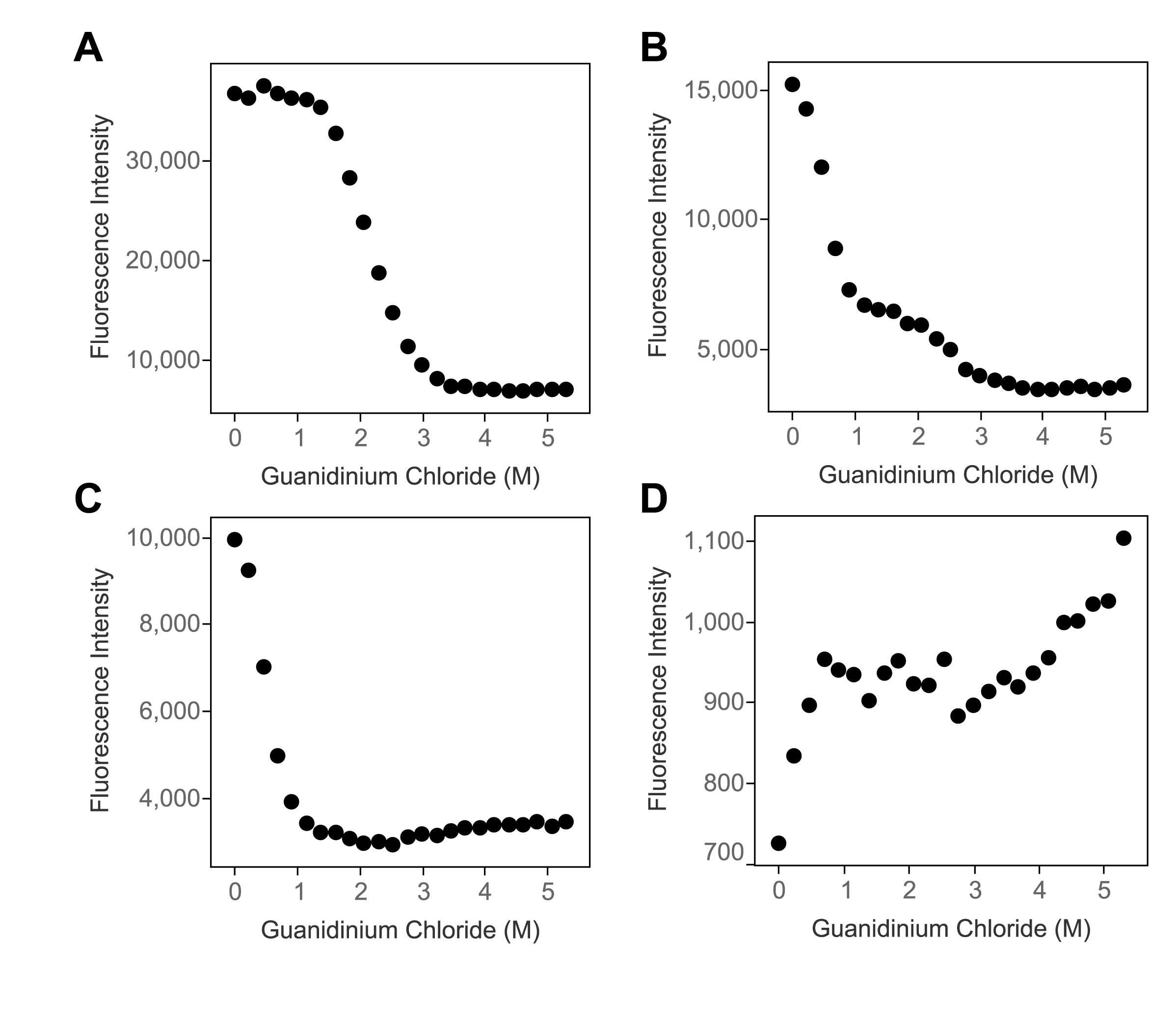

**Fig. S1.** Types of protein unfolding curves**.** Intrinsic Trp fluorescence in response to a gradient of the denaturant guanidinium chloride is used to determine protein stability by the linear extrapolation method (1). (*A*) WT Gβ1 features pre- and post-transition baselines flanking a smooth transition, all characteristics of a folded two-state protein that are required for stability determination. Certain mutations to Gβ1 can make accurate curve fitting untenable, as described in the next three panels. (*B*) The Y45Q variant exhibits two transitions and multiple baselines, which is indicative of a folding intermediate that invalidates the two-state assumption in the linear extrapolation method. (*C*) The A26M variant has no pre-transition baseline, indicating a folded, yet very destabilized protein preventing a quantitative stability measurement. (*D*) The A26R variant has a very low fluorescence signal and no observable transition, which are hallmarks of an unfolded or non-expressed protein.

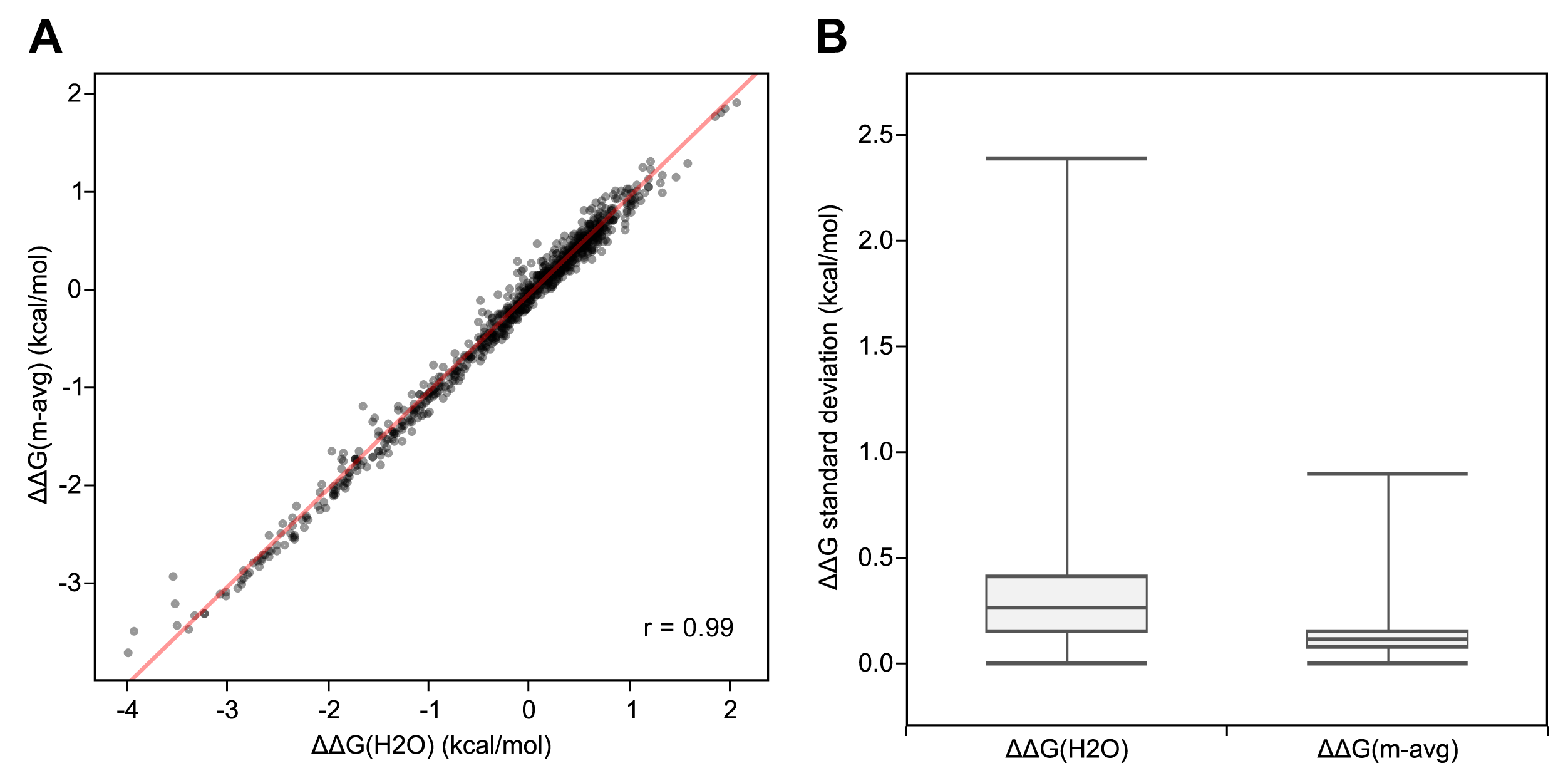

**Fig. S2.** Comparison of methods for calculating ΔΔ*G*. The average and standard deviation over multiple measurements of each single mutant is calculated for both ΔΔ*G* methods. (*A*) Mean ΔΔ*G*(H_2_O) plotted as a function of mean ΔΔ*G*(*m-*avg) exhibits a very strong linear relationship (slope = 0.99, *r* = 0.99) over the data set. The linear regression line is shown in red. (*B*) The distribution of standard deviations of ΔΔ*G* measurements for each calculation method. The ΔΔ*G*(*m-*avg) method (median = 0.10 kcal/mol) is more precise than ΔΔ*G*(H_2_O) (median = 0.26 kcal/mol), with very few extreme outliers.

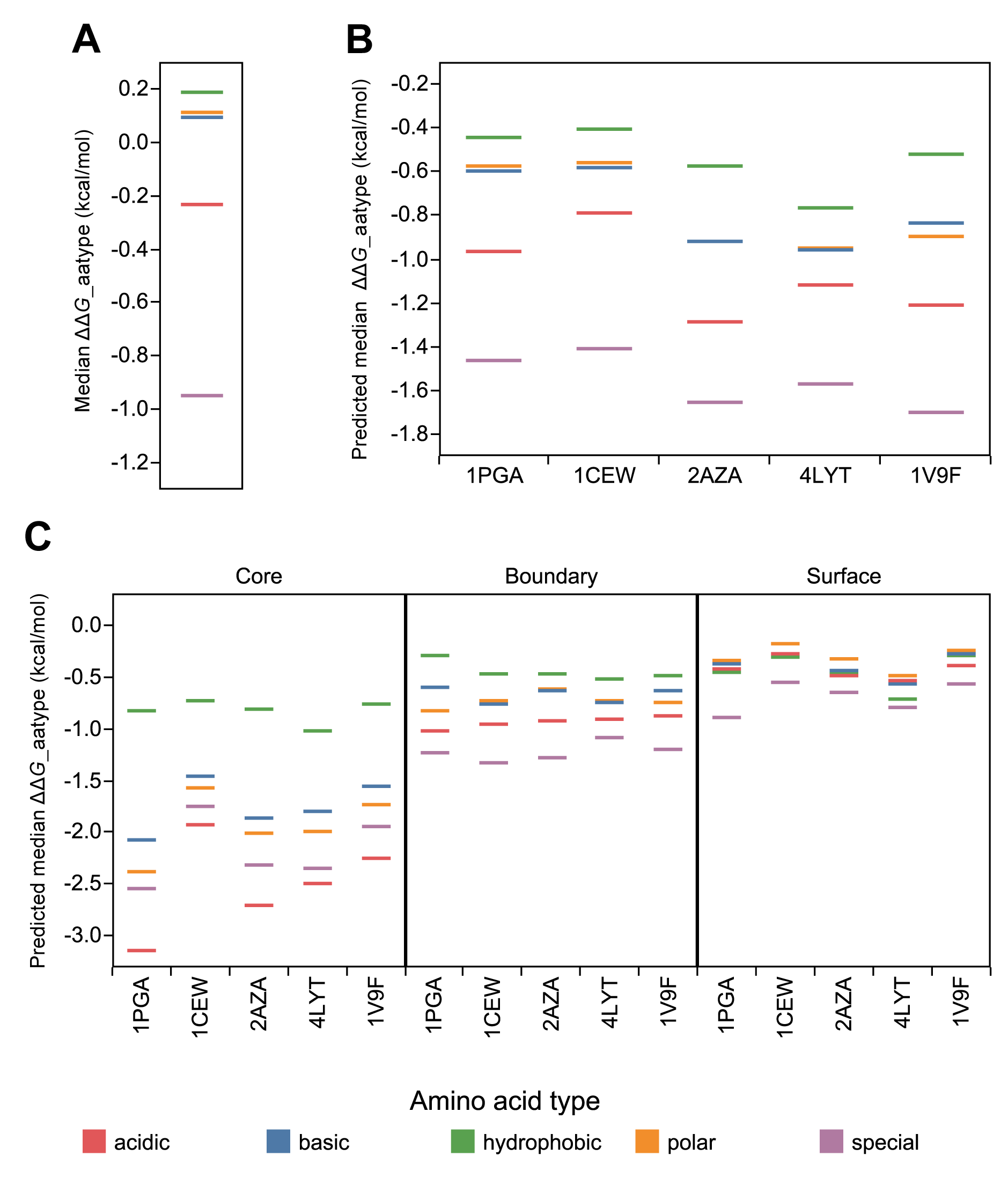

**Fig. S3.** Predicted ΔΔ*G* distributions grouped by mutant amino acid type recapitulate experimental trends. (*A*) Experimentally determined median ΔΔ*G* values by mutant amino acid type (aatype) for Gβ1 are plotted as Gantt lines. (*B*) Predicted median ΔΔ*G* values by mutant amino acid type are shown as Gantt lines for Gβ1, cystatin, azurin, lysozyme C, and pseudouridine synthase (PDB IDs: 1PGA, 1CEW, 2AZA, 4LYT, and 1V9R, respectively). Beyond Gβ1, the four other proteins were selected to span a range of sizes, secondary structure compositions, and packing densities, as these features could affect a protein's tolerance to different mutant amino acid types (Table S1). (*C*) Predicted median ΔΔ*G* values by mutant amino acid type are broken down by RESCLASS (4). Amino acids are colored by physiochemical type. Predicted ΔΔ*G* values were calculated using the PoPMuSiC 3 webserver (http://www.dezyme.com).

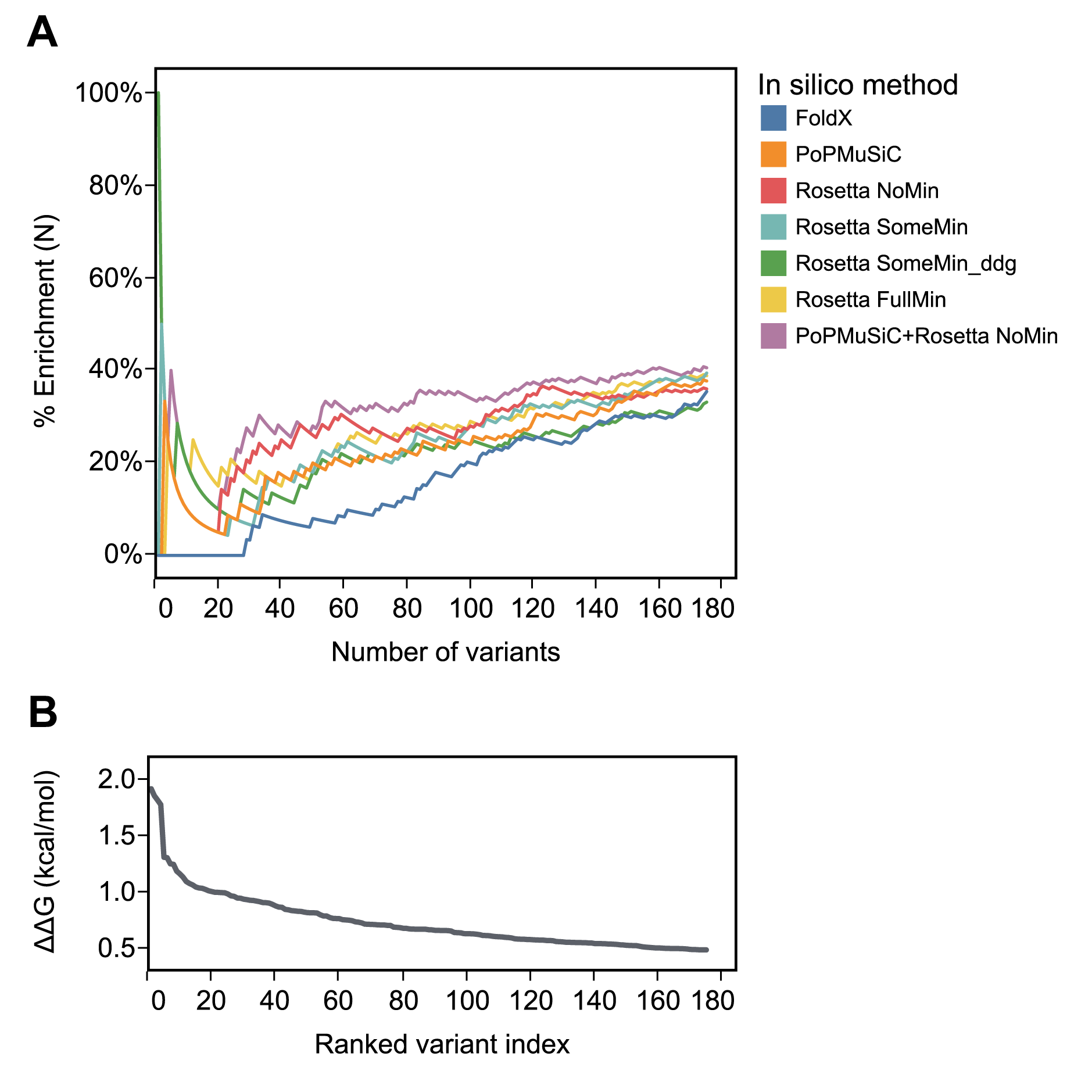

**Fig. S4.** Comparing stability prediction algorithms by % enrichment (%*E*). (*A*) %*E*(*N*), as defined in the text, is plotted as a function of the number of variants included in the list comparison. Only the top 175 Gβ1 single mutants are shown, sorted by ΔΔ*G*. (*B*) For reference, experimental ΔΔ*G* values are plotted as a function of the ranked variant index, a sorted list of the stability distribution. Each of the single algorithms and, for simplicity, only the best two-algorithm combination (PoPMuSiC+Rosetta NoMin) are shown, colored according to the legend.

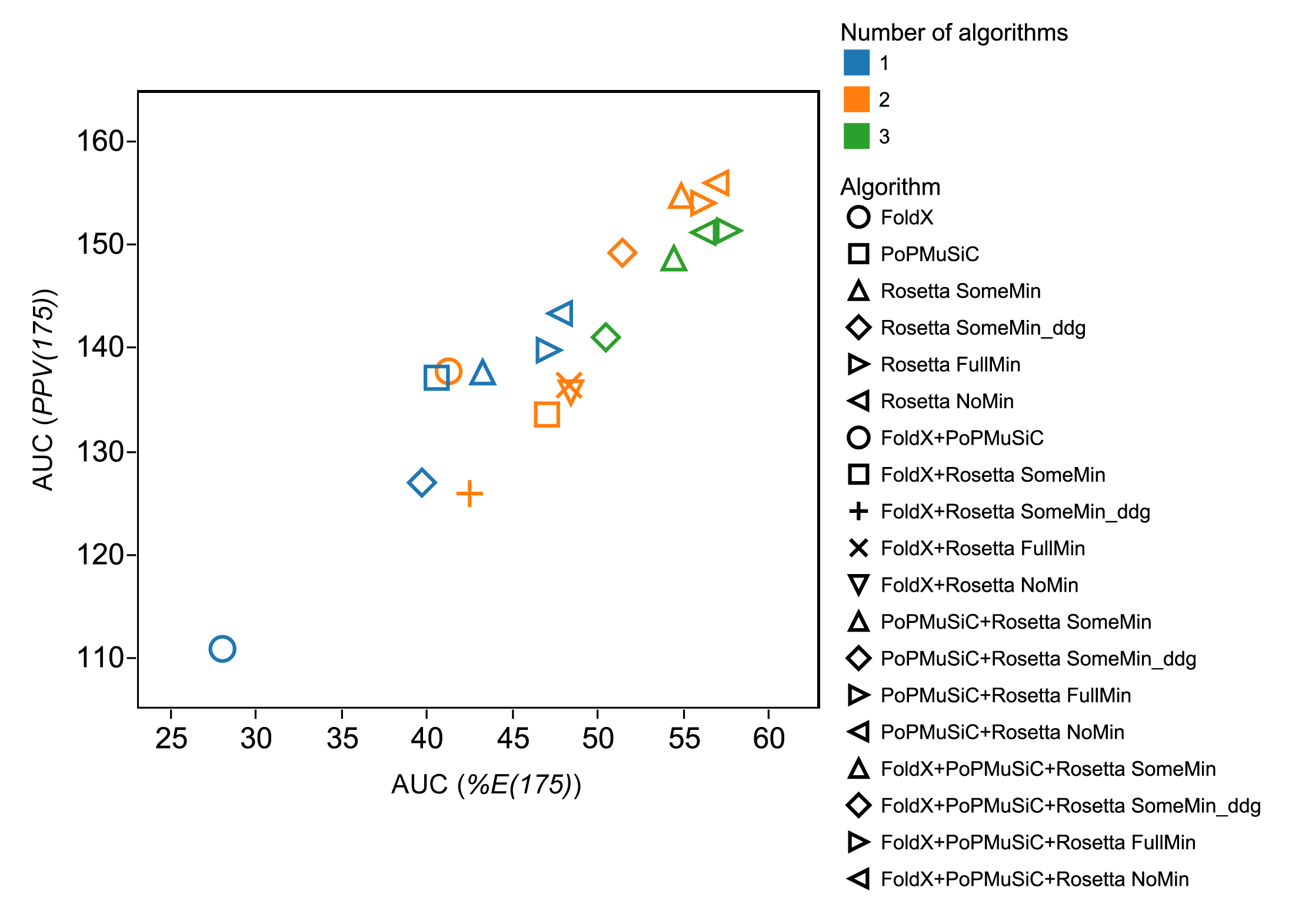

**Fig. S5.** Correlation between % enrichment (%*E*) and positive predictive value (*PPV*) area under the curve (AUC) values. The AUC is calculated for %*E* and *PPV* over the top 175 Gβ1 single mutants, sorted by ΔΔ*G.* Values for all single, double, and triple algorithm combinations are shown, colored and marked according to the legend.

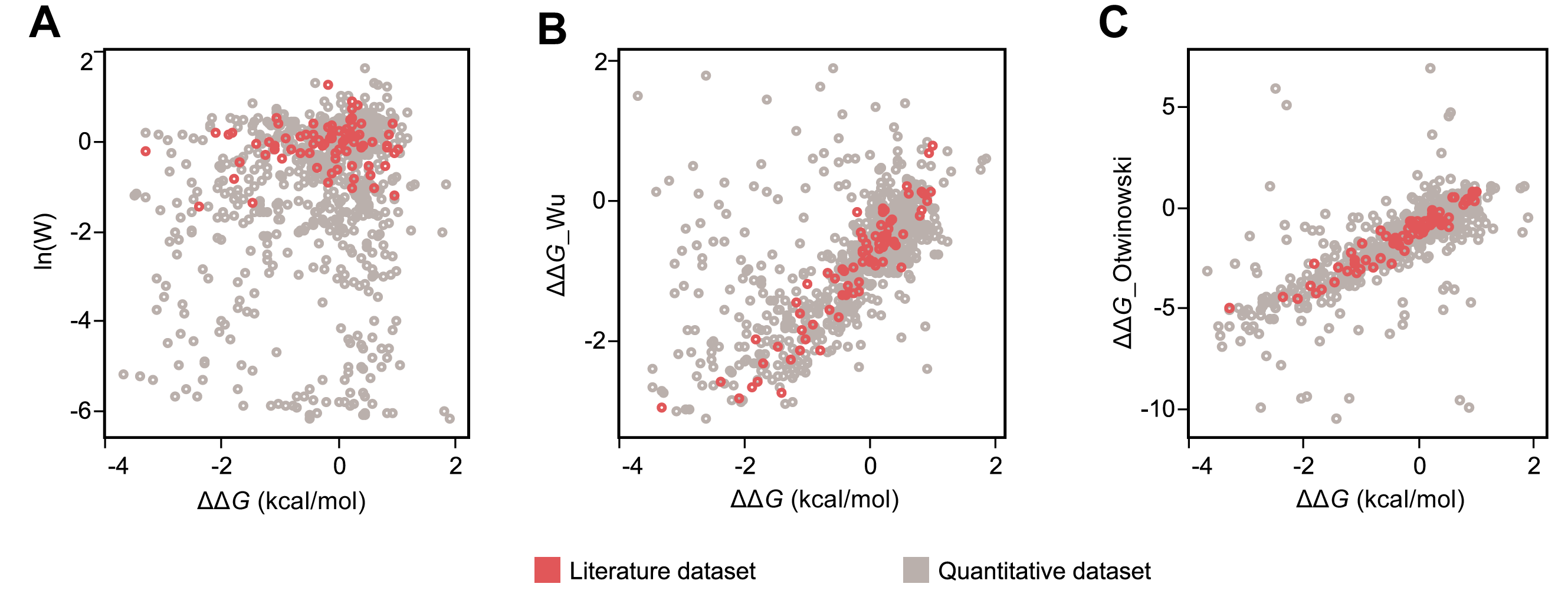

**Fig. S6.** Preexisting literature dataset used for validation does not accurately reflect deep mutational scanning (DMS) performance. Gβ1 single mutants common to the quantitative and depicted DMS datasets are shown. Red marks denote single mutants with corresponding data from the literature (∆*G*_lit) that were used to evaluate DMS performance. ∆∆*G*s from our experimental quantitative dataset (∆∆*G*) are plotted against (*A*) the natural log of fitness values, ln(W) obtained directly from DMS or (*B*) ∆∆*G* values predicted from the DMS data using the Wu et al. method (5) (∆∆*G_*Wu) or (*C*) *E*_folding values predicted from the DMS data using Otwinowski's three-state model (6) (∆∆*G_*Otwinowski). In all cases, DMS data is from Olson et al. (7).

**Table S1. Biophysical characteristics of selected proteins**

| PDB ID | Name | # of residues | % helix* | % strand* | Average OSP^†^ |
| --- | --- | --- | --- | --- | --- |
| 1PGA | Protein G1 | 56 | 25 | 42 | 0.333 |
| 1CEW | Cystatin | 108 | 20 | 48 | 0.321 |
| 2AZA | Azurin | 129 | 16 | 35 | 0.385 |
| 4LYT | Lysozyme C | 129 | 41 | 10 | 0.388 |
| 1V9F | Pseudouridine synthase | 249 | 20 | 22 | 0.383 |

*Secondary structure was determined by DSSP (8).

^†^Occluded surface packing (OSP) for each position is averaged over each protein. The difference in OSP for a position completely exposed on a loop and one involved in an alpha helix is ~0.25.

**Table S2. Algorithm performance by Spearman rank correlation**

|  |  | | |  | | Spearman correlation coefficient (*r*) | | | | | | | | | | |
| --- | --- | --- | --- | --- | --- | --- | --- | --- | --- | --- | --- | --- | --- | --- | --- | --- |
| Algorithm | | Backbone minimization* | | | Clash outliers^†^ | Overall | Surface**^‡^** | | Boundary**^‡^** | | Core**^‡^** | | +VolΔ | | −VolΔ | |
| PoPMuSiC | | |  | | 0 | 0.55 | | 0.55 | | 0.49 | | 0.38 | | 0.43 | | 0.70 |
| FoldX | | |  | | 17 | 0.46 | | 0.30 | | 0.57 | | 0.21 | | 0.30 | | 0.61 |
| Rosetta^§^ | | |  | |  |  | |  | |  | |  | |  | |  |
| NoMin | | | None | | 22 | 0.57 | | 0.43 | | 0.64 | | 0.21 | | 0.39 | | 0.79 |
| SomeMin | | | Constrained | | 17 | 0.58 | | 0.41 | | 0.66 | | 0.42 | | 0.38 | | 0.77 |
| SomeMin_ddg^¶^ | | | Constrained | | 6 | 0.55 | | 0.35 | | 0.62 | | 0.28 | | 0.35 | | 0.75 |
| FullMin | | | Unconstrained | | 3 | 0.56 | | 0.41 | | 0.63 | | 0.32 | | 0.37 | | 0.74 |

Predicted ΔΔ*G*s from stability algorithms were compared to experimental ΔΔ*G*s for Gβ1 single mutants in the quantitative dataset. Mutations with exceptionally high clash energies (clash outliers) were excluded when calculating each algorithm’s correlation coefficient. +VolΔ, small to large mutations; **−**VolΔ, large to small mutations.

*Level of backbone minimization after repacking for Rosetta methods.

^†^Number of mutations with a calculated repulsive energy > 2 standard deviations above the mean.

**^‡^**Residues are classified as core, boundary, or surface using RESCLASS (4).

^§^Rosetta parameter sets NoMin, SomeMin, and FullMin were initially described as row 3, row 16, and row 19, respectively (9).

^¶^Combines constrained backbone minimization with optimized reference energies trained on single mutant ΔΔ*G* data from ProTherm.

**Table S3. DMS performance by mutation class, volume change, and polarity change**

|  | Correlation coefficient (*r*) | |
| --- | --- | --- |
|  | ∆∆*G*_Wu | ∆∆*G*_Otwinowski |
| All mutations | 0.60 | 0.72 |
| Surface | 0.71 | 0.84 |
| Boundary | 0.63 | 0.82 |
| Core | 0.32 | 0.37 |
| +VolΔ | 0.55 | 0.64 |
| VolΔ | 0.60 | 0.78 |
| Nonpolar to polar | 0.37 | 0.61 |
| Nonpolar to nonpolar | 0.67 | 0.72 |
| Polar to polar | 0.63 | 0.74 |
| Polar to nonpolar | 0.72 | 0.83 |

DMS performance was measured as the correlation (*r*) between the experimental ∆∆*G* values for Gβ1 single mutants in our quantitative dataset and the predicted values obtained using the Wu et al. method (5) (∆∆*G*_Wu, *n* = 794) or the *E*_folding values predicted by Otwinowski (6) (∆∆*G*_Otwinowski, *n* = 812). In both cases, DMS data was from Olson et al. (7) Residues were classified as core, boundary, or surface using RESCLASS (4).

**References**

1.	Santoro MM, Bolen DW (1988) Unfolding free energy changes determined by the linear extrapolation method. 1. Unfolding of phenylmethanesulfonyl alpha-chymotrypsin using different denaturants. *Biochemistry* 27:8063-8068.

2.	Pace CN (1986) Determination and analysis of urea and guanidine hydrochloride denaturation curves. *Methods Enzymol* 131:266-280.

3.	Myers JK, Pace CN, Scholtz JM (1995) Denaturant m values and heat capacity changes: relation to changes in accessible surface areas of protein unfolding. *Protein Sci* 4:2138-2148.

4.	Dahiyat BI, Mayo SL (1997) De novo protein design: fully automated sequence selection. *Science* 278:82-87.

5.	Wu NC, Olson CA, Sun R (2015) High-throughput identification of protein mutant stability computed from a double mutant fitness landscape. *Protein Sci* 25:530-539.

6.	Otwinowski J (2018) Biophysical inference of epistasis and the effects of mutations on protein stability and function. *arXiv:* 1802.08744v2. Preprint, posted Mar 30 2018.

7.	Olson CA, Wu NC, Sun R (2014) A comprehensive biophysical description of pairwise epistasis throughout an entire protein domain. *Curr Biol* 24:2643-2651.

8.	Kabsch W, Sander C (1983) Dictionary of protein secondary structure: pattern recognition of hydrogen-bonded and geometrical features. *Biopolymers* 22:2577-2637.

9.	Kellogg EH, Leaver-Fay A, Baker D (2011) Role of conformational sampling in computing mutation-induced changes in protein structure and stability. *Proteins* 79:830-838.
